# Mitogen-activated protein kinases are carbon dioxide receptors in plants

**DOI:** 10.1101/2020.05.09.086116

**Authors:** Hanna Gałgańska, Łukasz Gałgański

**Affiliations:** Molecular Biology Techniques Laboratory, Faculty of Biology, Adam Mickiewicz University in Poznań, Uniwersytetu Poznańskiego 6, 61-614 Poznań, Poland

## Abstract

The amount of CO_2_ in the atmosphere is increasing continuously in the industrial era, posing a threat to the ecological balance on Earth. There are two ways to reduce elevated CO_2_ concentrations ([CO_2_]_high_): reducing human emissions or increasing their absorption by oceans and plants. However, in response to [CO_2_]_high_, plants diminish gas exchange and CO_2_ uptake by closing stomata. Surprisingly, we do not know how plants sense CO_2_ in their environment, and the basic mechanisms of the plant response to [CO_2_]_high_ are very poorly understood. Here, we show that mitogen-activated protein kinases (MAPKs) are plant CO_2_ receptors. We demonstrate that MPK4, a prominent MAPK that is known to be involved in the stomatal response to [CO_2_]_high_^1–3^, is capable of binding CO_2_ and is directly activated by a very low increase in [CO_2_] *in vivo* and *in vitro*. Unlike MPK4 activation by infections^4^, stress and hormones within known MAPK signalling cascades, [CO_2_]_high_-induced MPK4 activation is independent of the upstream regulators MKK1 and MKK2. Moreover, once activated, MPK4 is prone to inactivation by bicarbonate. The identification of stress-responsive MPK4 as a CO_2_ receptor sheds new light on the integration of various environmental signals in guard cells, setting up MPK4 as the main hub regulating CO_2_ availability for photosynthesis. This result could help to find new ways to increase CO_2_ uptake by plants.

## Introduction

Abscisic acid (ABA) is the best studied regulator of stomatal closure, and for many years, ABA-induced signalling events were thought to direct stomatal closure triggered by [CO_2_]_high_. Recent studies, however, have proposed otherwise, suggesting that both pathways work together and that ABA enhances the response to [CO_2_]_high_; however, [CO_2_]_high_ signalling is still active in the absence of ABA and key elements of ABA signalling^5^. Only the downstream effectors, S-type anion channels, i.e., SLAC1, and some of their regulators are shared by the ABA and CO_2_ pathways. Thus, except BIG^6^ and RHC1^7^, the connections of which with core pathways remain unclear, known specific regulators of CO_2_ signalling in guard cells belong to the MAPK superfamily. Among these proteins, CBC1/2^8^ and HT1 mitogen-activated protein kinase kinase kinases (MKKKs) are involved in pathways leading to low [CO_2_]-induced stomatal opening or inhibition of stomatal closure rather than [CO_2_]_high_-induced stomatal closure. In contrast, MPK12 and MPK4 are essential upstream mediators of the [CO_2_]_high_^1,2^ pathway promoting SLAC1-mediated stomatal closure via HT1^3^ inactivation.

Because MPK12 orthologues are guard cell-specific kinases found only in *Brassicaceae*^9^, we focused on MPK4 to reveal the general mechanisms of [CO_2_]_high_ sensing in all plants, as silencing of *Nt*MPK4 impaired [CO_2_]_high_- and dark-induced stomatal closure by disrupting the activation of slow-type anion channels in *Nicotiana tabacum*^1^.

## Results

### MPK4 is activated by CO_2_ *in vivo*

No kinase involved in plant CO_2_ signalling has been shown to be activated by [CO_2_]_high_ *in vivo* to date; therefore, we decided to study MPK4 activation in response to [CO_2_]_high_ in epidermal peels. In line with the reported high activity of the *MPK4* promoter in guard cells^4^, we detected very high MPK4 expression in Arabidopsis epidermal peels using immunoblotting (Fig. 1a).

**Fig. 1.**
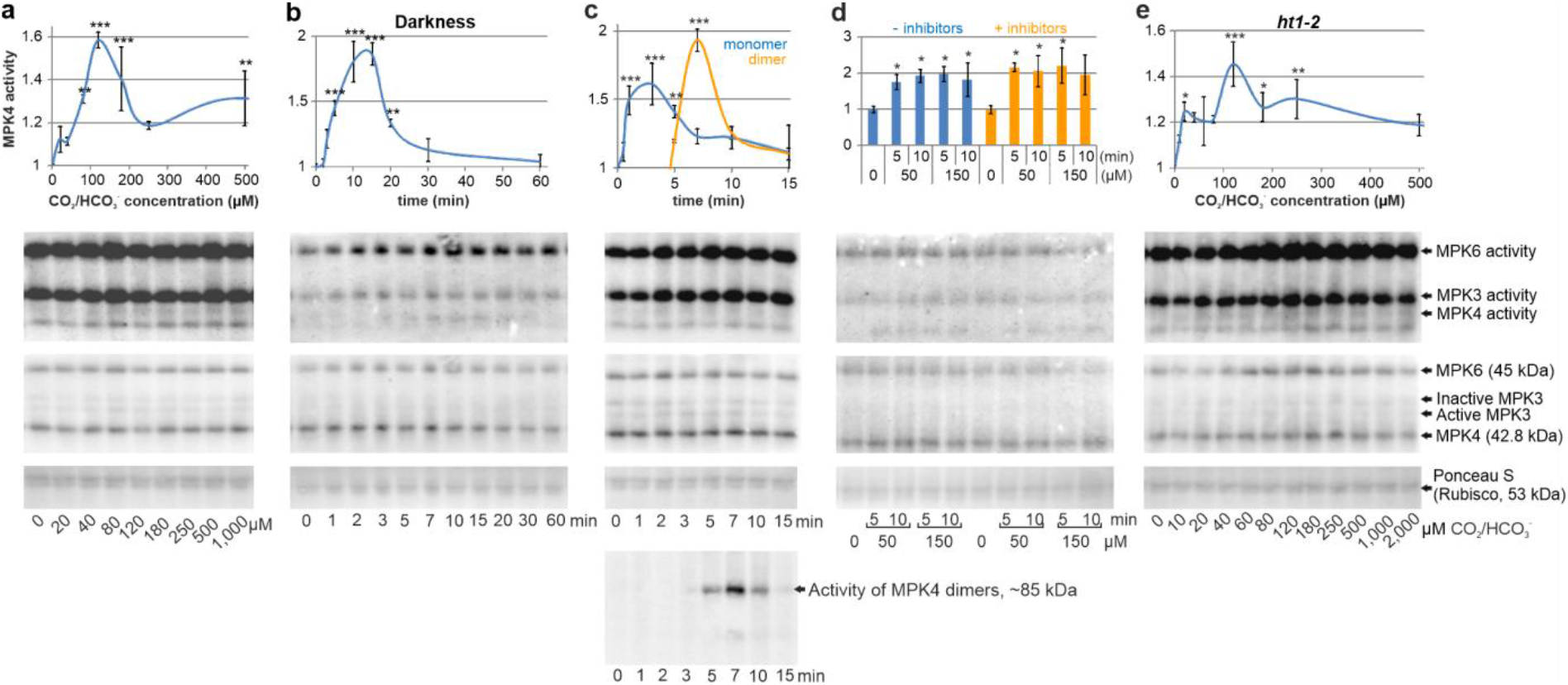
Activation of MPK4 by [CO_2_]_high_ in Arabidopsis epidermal peels. **a**, Measurement of MPK4 activity in response to the indicated [CO_2_]. **b, c**, Time course of MPK4 activation by darkness (**b**) and 180 μM HCO_3_^-^ (**c**). Strong activation of 85-kDa MPK was identified in response to 7-minute exposure to HCO_3_^-^ (orange line). Identical molecular mass, appearance time points and intensity changes in the 85-kDa protein band were found for both anti-TEY and anti-MPK4 (Supplementary Fig. 2) antibodies, indicating that dimeric MPK4 is ~85-kDa active MPK. **d**, MPK4 activation by [CO_2_]_high_ is independent of MAPK cascades – MKK inhibitors do not abolish MPK4 activation by [CO_2_]_high_. Epidermal peels were preincubated with both 50 μM PD98059 and 5 μM U0126 for 1.5 h before addition of the indicated concentration of dissolved CO_2_. **e**, MPK response to [CO_2_]_high_ in *ht1-2*. MPK4 activity was studied in an open system: epidermal peels were incubated in stomatal opening buffer in open tubes, ensuring continuous CO_2_ exchange with ambient air. Then, darkness or specified CO_2_ concentrations were applied for the indicated time or 15 min, respectively. To gain insight into MPK4 activity and separate it from highly active MPK3, high-resolution electrophoresis was applied, followed by immunodetection of active MPKs with phospho-p44/42 MAPK (Erk1/2) (Thr202/Tyr204) antibody (anti-phospho-TEY) against the phosphorylated activation loop of MPKs. Protein loading was visualized by both Ponceau S staining and immunoblotting with anti-MPK3, anti-MPK4 and anti-MPK6 antibodies. Representative results from three independent experiments are presented. Error bars represent the standard deviation (SD). *, ** and *** indicate significant differences in MPK4 activity (p<0.05, p<0.01 and p<0.001, respectively) compared to the control. The above data were obtained on proteins isolated by phenol-SDS extraction for immediate separation of ATP^33^ from MPKs to prevent their extracellular activation. In contrast, we were unable to detect the activity of guard cell MPK4 purified under native conditions (Supplementary Fig. 9).

The activity of MPK4 was extremely low compared to that of the highly active MPK3 and MPK6. The lack of MPK4 activation in the control samples indicates the maintenance of stress-free conditions in the experimental system used. We assumed that the method used to study [CO_2_]_high_-induced MPK activation should utilize direct analysis of the protein extract without lengthy sample preparation steps at indoor [CO_2_] under native conditions. Therefore, we rejected the classic in-gel kinase assay following kinase immunoprecipitation.

MPK4 is activated by as low as 20 μM CO_2_/HCO_3_^-^, reaching the highest activity in 120 μM CO_2_/HCO_3_^-^. Consistent results were obtained when the source of CO_2_/HCO_3_^-^ was dissolved CO_2_ (Fig. 1a) or KHCO_3_ (Supplementary Fig. 1a-b). As stomatal closing in darkness is a typical physiological response to [CO_2_]_high_ and is independent of arbitrarily imposed external CO_2_ concentrations, we traced the effect of darkness on MPK4 activity over time. The result revealed MPK4 activation by darkness at time points from 5 to 20 min (peak at 10-15 min, p<0.001, Fig. 1b) compared to immediate (maximum at 2-5 min) activation by externally provided [CO_2_]_high_ (Fig. 1c) according to the very rapid stomatal closure in response to [CO_2_]_high_^10^. A decrease in monomeric MPK4 activity was accompanied by strong (p<0.0001) and transitory activation of the ~85-kDa form of MPK4 (Fig. 1c, Supplementary Fig. 2) corresponding to the MPK4 dimer in size according to the multimerization of active MPK4 in response to both CO_2_ (Galganska et al., in preparation) and H_2_O_2_^11^.

Generally, MAPKs function as a cascade in which MKKK phosphorylates and activates a mitogen-activated protein kinase kinase (MKK), which in turn activates an MPK. Therefore, a lack of [CO_2_]_high_-induced MPK4 activation would be expected in plants with blocked upstream MKKs if activation of MPK4 was a part of the secondary response to [CO_2_]_high_ within the MAPK cascade. Thus, we measured MPK4 activity in epidermal peels pre-treated with MKK inhibitors (PD98059 and U0126; Fig. 1d) and found that the increase in MPK4 activity in response to [CO_2_]_high_ was still statistically significant, indicating MAPK cascade-independent activation of MPK4. Furthermore, we found that MPK4 activation in response to [CO_2_]_high_ was intact in *ht1-2* (Fig. 1e), supporting previous data^3^ showing the MPK4 position upstream of HT1.

### MPK4 is activated by CO_2_ *in vitro*

Based on the response of MPK4 to [CO_2_]_high_ independent of upstream signalling (i), the upstream role of this protein in known CO_2_ signalling components (ii) and its importance in CO_2_ signalling (iii), we hypothesized that the role of MPK4 is that of a direct CO_2_ sensor. Thus, we measured MPK4 activity in response to [CO_2_]_high_ *in vitro*.

A CO_2_ receptor is expected to sense very low [CO_2_] because guard cells are able to react to slight changes in ambient [CO_2_], and the dissolved atmospheric [CO_2_] in the acidic pH of the apoplast is expected to be slightly above 10 μM. Moreover, upon its transport through the cell membrane, CO_2_ is spontaneously converted to HCO_3_^-^ at cytoplasmic pH and further consumed by photosynthesis. There is no report clearly showing [CO_2_] in guard cells. In addition, net CO_2_ uptake or production from mitochondrial respiration, photorespiration and photosynthetic CO_2_ fixation remains unclear in guard cells. Typically, the intracellular partial pressure of carbon dioxide (pCO_2_) in photosynthetic cells reaches approximately half the pCO_2_ concentration in the ambient air, but CO_2_ is unequally distributed within the cell^12^.

Based on these assumptions, MPK4 is activated by as low as 5 μM dissolved CO_2_ (Fig. 2a) or KHCO_3_ (Supplementary Fig. 3a-b) added to the *in vitro* phosphorylation mixture. MPK4 activation occurs in just a few seconds (Supplementary Fig. 3c), as shown by an increase in activation loop autophosphorylation. An increase in substrate protein phosphorylation by MPK4 was observed 3 min after CO_2_ administration (Fig. 2b). To carefully exclude any artefacts, we investigated MPK4 activation in several systems using GST-tagged MPK4 (Fig. 2b, Supplementary Fig. 3c) and tag-free MPK4 (Fig. 2a, Supplementary Fig. 3a) dephosphorylated by FastAP alkaline phosphatase. We used an anti-phospho-TEY antibody (Fig. 2a, Supplementary Fig. 3c) or an *in vitro* kinase assay using commercial myelin basic protein (MBP) as a standard MPK substrate (Supplementary Fig. 3b,d) or recombinant JAZ12 as a specific and natural substrate of MPK4 (Fig. 2b,c). All of the abovementioned approaches confirmed that [CO_2_]_high_ promoted MPK4 activation. However, only a low increase in [CO_2_] influenced MPK4 activity with a constant trend in the *in vitro* kinase activity assay; the application of 40 μM CO_2_ or higher yielded variable results (Supplementary Fig. 3d), suggesting that MPK4 activity can be affected by both CO_2_ forms, namely, free CO_2_ and HCO_3_^-^, with opposite effects. Thus, we measured [CO_2_]_high_-induced MPK4 activation in a pH-dependent manner because at low pH, the CO_2_/HCO_3_^-^ equilibrium is shifted towards increased free [CO_2_], whereas at high pH, the equilibrium is shifted towards increased [HCO_3_^-^]. An increase in [CO_2_/HCO_3_^-^] clearly activates MPK4 at low pH in contrast to high pH (Fig. 2c), indicating that an increase in [CO_2_] enhances MPK4 kinase activity and an increase in [HCO_3_^-^] reduces MPK4 kinase activity. The negative effect of HCO_3_^-^ on MPK4 activity was further confirmed in experiments with constant [CO_2_] and increasing [HCO_3_^-^] (Supplementary Fig. 4a) and by direct comparison of [CO_2_] and [HCO_3_^-^] (Supplementary Fig. 4b).

**Fig. 2.**
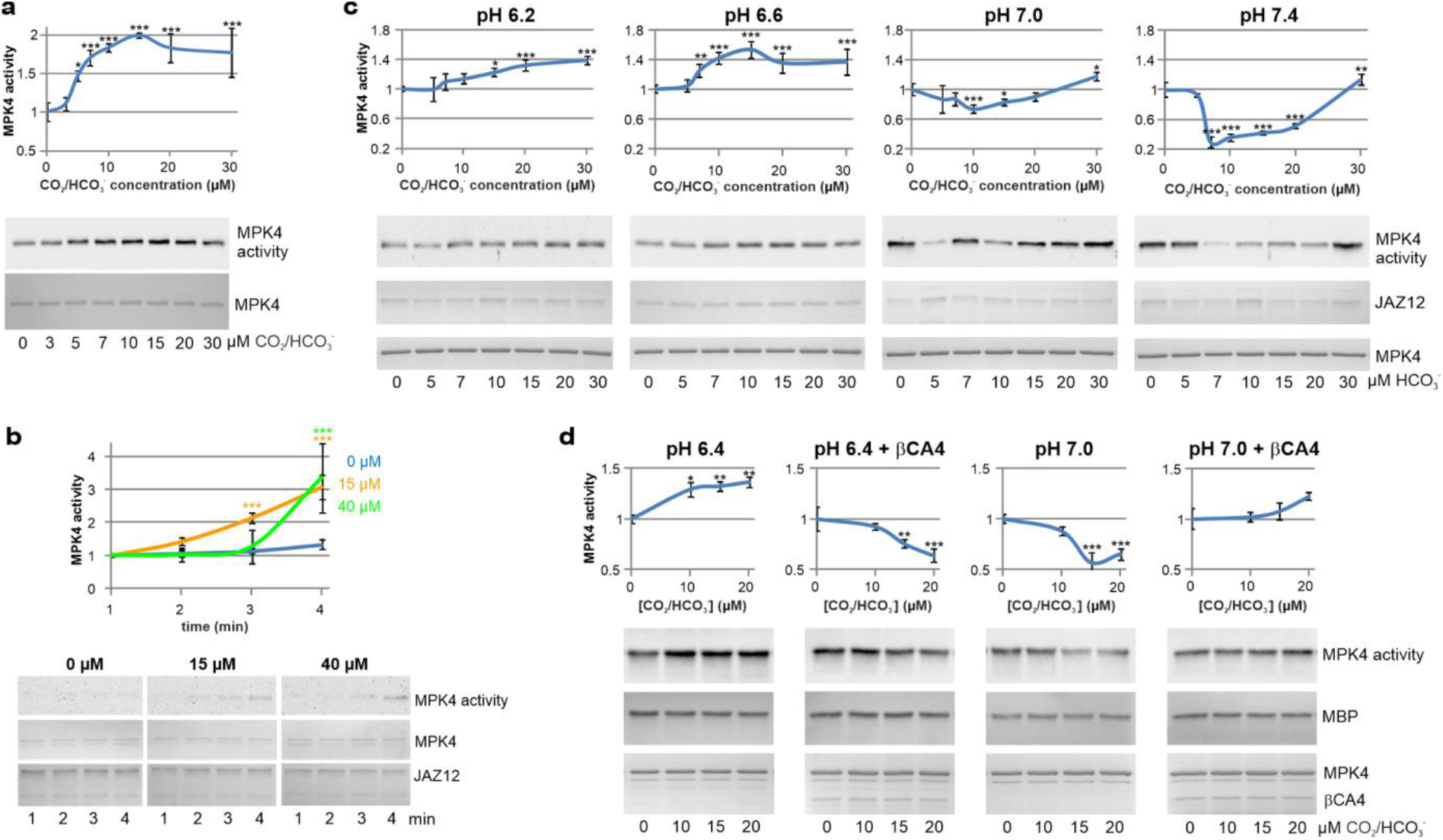
[CO_2_]_high_ directly activates MPK4 *in vitro*. **a**, [CO_2_]_high_ enhances the phosphorylation of the MPK4 kinase activation loop, as shown by immunoblotting using anti-phospho-TEY. The *in vitro* phosphorylation reaction at pH 7.0 was carried out at 24°C for 1 min upon the addition of the indicated [CO_2_]. **b**, Time course of MPK4 activation by HCO_3_^-^ at pH 6.4 presented as the intensity of JAZ12 thiophosphorylation using immunoblotting with anti-thiophosphate ester antibody (anti-TE). **c**, pH-dependent MPK4 activity regulation by HCO_3_^-^. Thiophosphorylation (24°C, 15 min) of JAZ12 followed by immunoblotting with anti-TE. **d**, HCO_3_^-^/CO_2_ conversion by βCA4 reverses the pH-dependent MPK4 activation pattern. *In vitro* phosphorylation reactions with MBP as a substrate were preincubated for 25 min in the presence or absence of βCA4. Then, MPK4 was added and incubated for 25 min. MPK4 activity was detected by immunoblotting with anti-phospho-MBP. Experiments in (**a**-**d**) were carried out using MPK4 purified from bacteria and dephosphorylated by FastAP phosphatase GST-MPK4 (**b**, **c**) or tag-free MPK4 (**a**, **d**). Quantities of substrate proteins (**b**-**d**) and MPK4 (**a**) were visualized by Ponceau S staining. βCA4 and MPK4 in **b**-**d** were stained with Coomassie Brilliant Blue R-250 (CBB). Representative results from three independent experiments are presented; mean +SD; *, ** and *** indicate p<0.05, p<0.01 and p<0.001, respectively.

One could wonder how MPK4 functions in cells, where the pH of the cytoplasm (7.0-7.2) promotes HCO_3_^-^ formation. MPK4 could be activated *in vivo* due to the action of carbonic anhydrases (CAs), which were shown to be essential for the CO_2_ signalling pathway^13^. As CAs act in both directions to regulate the CO_2_/HCO_3_^-^ equilibrium, we added βCA4, one of the two most abundant Arabidopsis CAs^13,14^, to *in vitro* phosphorylation reactions. At pH 7.0, βCA4 increased [CO_2_] and reversed the MPK4 activity profile from MPK4 inactivation to MPK4 activation. Consequently, at pH 6.4, βCA4 increased [HCO_3_^-^], leading to MPK4 inactivation instead of activation in the absence of βCA4 (Fig. 2d). These results support the positive role of CO_2_ and the negative role of HCO_3_^-^ in MPK4 activation and demonstrate that the CO_2_/HCO_3_^-^ equilibrium, not pH, regulates MPK4 activity.

### MPK4 binds CO_2_

The CO_2_ receptor is expected to bind CO_2_. We verified that MPK4 efficiently bound ^14^CO_2_ (p<0.001) compared to both BSA and the sample devoid of protein (Fig. 3a, Supplementary Fig. 5a). However, it was not possible to precisely determine the K_D_ for the MPK4-^14^CO_2_ interaction because the experiments were carried out in an open system with free exchange of diluted ^14^CO_2_ with ambient atmosphere (i), ^12^CO_2_ was also available for MPK4 (ii), and possible competitive binding of H^14^CO_3_^-^ to MPK4 (iii). However, ^14^CO_2_ binding by MPK4 at low pH showed two maxima, and the first peak was reached at 10 μM ^14^CO_2_/H^14^CO_3_^-^ (4.88 μM and 3.76 μM ^14^CO_2_ at pH 6.4 and 6.6, respectively; Fig. 3b) when the molar ratio of ^14^CO_2_ and MPK4 was approximately 1:1. Importantly, the graphs of ^14^CO_2_ binding with increasing [^14^CO_2_] in the pH series closely reflect the MPK4 activity graphs under the same conditions (Supplementary Fig. 5b-d). The coincident decrease in both MPK4 activity and ^14^CO_2_ binding (in 15-20 μM CO_2_/HCO_3_^-^ at pH 6.4 and 6.6) indicates the stronger binding of ^14^CO_2_ than that of H^14^CO_3_^-^. Taken together, the above data support the designation of MPK4 as a CO_2_ receptor.

**Fig. 3.**
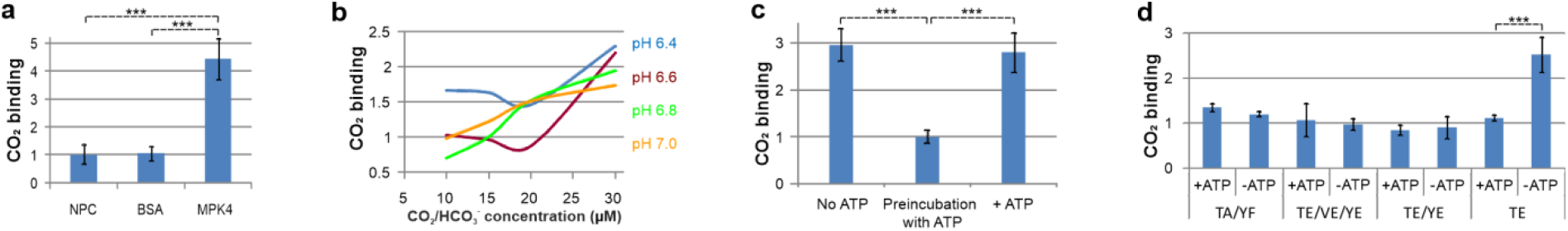
MPK4 binds CO_2_. **a**, ^14^CO_2_ (10 μM H^14^CO_3_^-^ incubated at pH 6.4) is effectively coeluted from a size-exclusion chromatography column with MPK4 in contrast to BSA or a no-protein control (NPC). The experimental design is illustrated in Supplementary Fig. 5a. **b**, Efficiency of CO_2_ binding at increasing [^14^CO_2_] in a pH series. Charts of individual pH series, including error bars, are shown in Supplementary Fig. 5b. **c**, MPK4 autophosphorylation prevents effective CO_2_ binding. MPK4 preincubated with ATP for 2 min is not able to bind CO_2_ in contrast to both reactions containing ATP without a preincubation step or ATP-free reactions. **d**, MPK4 with only T201 of TEY phosphorylated is still able to bind CO_2_ in the absence of ATP, in contrast to MPK4 with double TEY phosphorylation. In **c**-**d**, MPK4 incubation with 15 μM H^14^CO_3_^-^ was carried out for 10 min at pH 7.0. Plotted values of disintegrations per minute (DPM) after normalization based on background radioactivity in individual experiments. Mean ±SD; n=3; *** indicates p<0.001.

### Active MPK4 is prone to inactivation by HCO_3_^-^

To obtain further insight into the opposing effects of HCO_3_^-^ and CO_2_ on MPK activity, we measured the [CO_2_]_hig_h-induced activation of several MPKs at pH 7.0 (Fig. 4a-b). It turned out that the higher MPK activity was under control conditions, the stronger the HCO_3_^-^-induced inactivation of MPKs, and MPKs with low basal kinase activity (MPK12, MPK20, and *Hv*MPK4) were activated in response to [CO_2_]_hig_h without the effect of kinase inactivation. Because MPK activity depends on the phosphorylation of conserved TEY or TDY motif in the kinase activation loop, we investigated the impact of TEY phosphorylation on MPK4 activity regulation by both CO_2_ and HCO_3_^-^ using MPK4 versions with modified TEY motif (Fig. 4c-e).

**Fig. 4.**
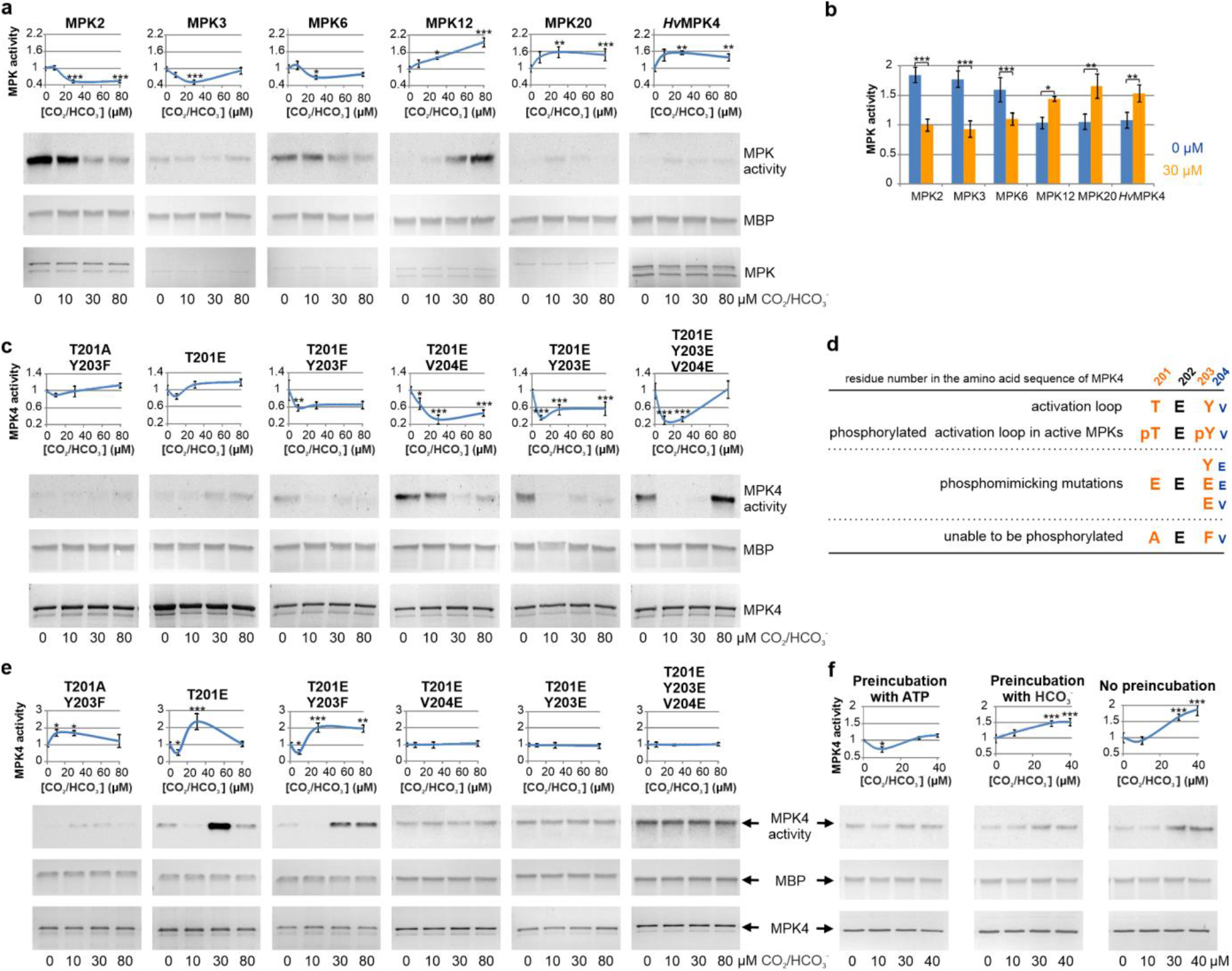
The MPK response to CO_2_ is governed by initial MPK activity. **a**, Highly active MPKs are downregulated by HCO_3_^-^ in contrast to MPKs with low kinase activity. The MPK4 homologue from barley was included in this analysis due to the quickest response of barley stomata to darkness among the studied species^34^. For more details, see Supplementary Fig. 10. GST-MPK fusion proteins for *in vitro* phosphorylation were purified from bacteria. **b**, A summary of the data presented in **a**. **c**, Mutations presented in **d** alter the MPK4 response to [CO_2_]_high_. **d**, Schematic representation of generated mutations mimicking phosphorylated or unphosphorylable amino acids in the kinase activation loop (red) and the following valine (blue) in MPK4. **e**, Increasing the CO_2_/HCO_3_^-^ ratio by lowering the pH from 7.0 (shown in **c**) to 6.6 disables the inhibition of active versions of MPK4 by HCO_3_^-^ and indicates that T201 is responsible for this effect. **f**, WT MPK4 inactivation by 10 μM HCO_3_^-^ is promoted by MPK4 phosphorylation. Preincubation with either HCO_3_^-^ or ATP was carried out for 10 min (24°C). Dephosphorylated MBP was used as an MPK4 substrate during *in vitro* phosphorylation (30 min, 30°C), followed by immunoblotting with anti-phospho-MBP. In **a**, **c**, **e**, kinase activity was measured by *in vitro* MBP thiophosphorylation (15 min, 24°C) detected by immunoblotting using anti-TE. Dephosphorylated kinases were used in all assays to exclude any effects of phosphorylated amino acids other than the TEY motif. Loading of MPKs was visualized by CBB, while MBP was visualized by Ponceau S. Representative results from three independent experiments are presented. Mean ±SD; *, ** and *** indicate p<0.05, p<0.01 and p<0.001, respectively.

Mutants mimicking MPK4 with phosphorylated Y203 of TEY (MPK4^T201E/Y203E^, MPK4^T201E/V204E^ and MPK4^T201E/Y203E/V204E^), reflecting full MPK4 activity, could not be further activated by [CO_2_]_high_, whereas unphosphorylated MPK4 (MPK4^T201A/Y203F^) and MPK4 with only T201 phosphorylated (MPK4^T201E^) were still prone to [CO_2_]_high_-induced activation (Fig. 4c). Thus, [CO_2_]_high_ not only promotes TEY phosphorylation (Fig. 2a) but also acts as an additional activity enhancer of inactive or incompletely activated MPK4.

All the mutants tested were negatively regulated by HCO_3_^-^ at pH 7.0 (~85% HCO_3_^-^ and ~15% CO_2_) (Fig. 4c). Lowering the pH to 6.6 (~62% HCO_3_^-^ and ~38% CO_2_) eliminated HCO_3_^-^-induced inactivation of all MPK4 forms with phosphorylated Y203 (Fig. 4e). This result is consistent with the decrease in ^14^CO_2_/H^14^CO_3_^-^ binding in the concentration range of 15-20 μM at pH 6.6 (Fig. 3b) and further supports lower binding of HCO_3_^-^ than of CO_2_ by MPK4. Importantly, T201 phosphorylation (MPK4^T201E^, MPK4^T201E/Y203F^), in contrast to unphosphorylated T201 (MPK4^T201A/Y203F^), enhances MPK4 susceptibility to inhibition by HCO_3_^-^ (Fig. 4e).

The effects of TEY phosphorylation on HCO_3_^-^-triggered inhibition of MPK4 were further confirmed using WT MPK4. MPK4 preincubated with ATP (autophosphorylated on TEY) before CO_2_ addition is prone to strong HCO_3_^-^-induced inactivation, in contrast to preincubation of MPK4 with CO_2_ before ATP application or administration of both ATP and CO_2_ at the same time (Fig. 4f). This indicates competition between CO_2_ and HCO_3_^-^. The effect of HCO_3_^-^ becomes noticeable at high [HCO_3_^-^] or at pH ≥7 (high [HCO_3_^-^]/[CO_2_] ratio) only when the TEY of MPK4 is already phosphorylated. Accordingly, HCO_3_^-^ does not influence [CO_2_]_high_-induced TEY phosphorylation; in contrast to the decrease in MPK4 activity observed as MBP or JAZ12 phosphorylation at pH ≥7, [CO_2_]h_ig_h-induced TEY phosphorylation is not inhibited by HCO_3_^-^ (Supplementary Fig. 6).

As studies on kinase activity can be conducted only in the presence of ATP, we employed a ^14^CO_2_ binding assay to further investigate the role of ATP in CO_2_ sensing by MPK4. MPK4 preincubation with ATP impaired ^14^CO_2_ binding 3-fold compared to that observed when both ^14^CO_2_ and ATP were added at the same time or when ^14^CO_2_ preincubation was conducted before ATP delivery (Fig. 3c). However, ATP does not influence the ^14^CO_2_ binding of MPK4 mutants mimicking phosphorylated TEY (MPK4^T201E/Y203E^, MPK4^T201E/Y203E/V204E^). In contrast, ATP diminished (3-fold) ^14^CO_2_/H^14^CO_3_^-^ binding by MPK4^T201E^ (Fig. 3d), indicating that the transition from pTEY to pTEpY is crucial for ATP-dependent ^14^CO_2_/H^14^CO_3_^-^ binding. Moreover, the weakened CO_2_ binding by MPK4^T201A/Y203F^ supports the importance of TEY for CO_2_ recognition.

The lack of CO_2_ binding under high ATP availability may underlie the mechanism for elimination of fluctuations in endogenous [CO_2_], because when ATP availability increases, [CO_2_] increases locally due to the proximity of mitochondria. Interestingly, MPK4 inactivation is strongest at concentrations of dissolved atmospheric CO_2_ and at very high [CO_2_]. Physiologically, such a strong HCO_3_^-^-induced inhibitory effect on activated MPK4 seems to be a very effective autoregulatory mechanism, in which the CO_2_ sensor is inactivated during a long-term increase in [CO_2_] (mainly HCO_3_^-^ at the pH of the cytoplasm). This may also be an important mechanism of cross-talk between CO_2_ and stress signalling, as different adverse conditions activate MPK4, leading to modification of the plant response to CO_2_ during stress (Supplementary Fig. 7).

In general, our findings are important for the regulation of plant growth and development by CO_2_, as MPK4 regulates cytokinesis^15^ and photosynthesis^16^. The best summary of this is a picture of the highly enlarged stomata (Supplementary Fig. 8) of extremely dwarfed *mpk4* plants^4,16^, supporting previous results from tobacco plants with silenced *Nt*MPK4^1^.

The broad importance of the presented results could be considered because MAPKs are conserved enzymes in all eukaryotes. In human lungs, MAPKs are activated by SARS-CoV, SARS-CoV-2^17,18^ and other causative agents of pneumonia^19–23^ to trigger the production of proinflammatory cytokines. Angiotensin-converting enzyme 2 (ACE2) inhibits MAPK signalling^19^ and thus protects against severe lung diseases caused by lipopolysaccharide^20,21^, bleomycin^19^, and cigarette smoke^22^ and particulate matter 2.5 (PM2.5) exposure^23^. However, ACE2 is bound by SARS-CoV-2^24,25^, leading to cytokine storms and a severe course of pneumonia and resulting in acute respiratory distress syndrome (ARDS) and pulmonary fibrosis. Therefore, the inhibition of active MAPKs could be a strategy to prevent the acute course of COVID-19. Based on the inactivation of active plant MPKs by CO_2_ described herein, we encourage researchers to study the inhibitory effect of CO_2_ on human MAPKs because both synthetic MAPK inhibitors^26^ and ten-minute inhalation of 5% CO_2_^27^ protect against lipopolysaccharide-induced lung injury in mice. In addition, tobacco smoke has been suggested recently to be a protective factor against the development of COVID-19 symptoms. Importantly, CO_2_ is a natural and safe gas in the lungs, and short-term CO_2_ inhalation is beneficial for the respiratory, nervous^28–30^ and circulatory^31,32^ systems.

## Methods

### General considerations

All protein purifications, handling of purified proteins and experiments using extracted proteins were carried out in empty rooms (max. 2 persons/40 m^2^) with open windows providing fresh air. During the heating season, no research was conducted on windless days or when the PM10 concentration in air exceeded 30 μg m^-3^. The breath was not directed towards the open tubes and pipette tips. Ice was not used due to the reduction in CO_2_ solubility with increasing temperature and because of ice production from high-pH water in our laboratory. All solutions were prepared using acidified (pH 4.8-5.2) CO_2_-free water in rooms with fresh air. Solutions were stored frozen, or the pH was adjusted immediately before use. MPK purification or modification (e.g., dephosphorylation or protease digestion) was followed by protein desalting using Amicon Ultra filters (Millipore, Billerica, MA) to remove HCO_3_^-^ and other salts and buffers.

Solutions containing the indicated CO_2_ or HCO_3_^-^ concentrations were prepared from freshly dissolved 100 mM KHCO_3_^-^ or CO_2_-saturated water. The CO_2_ concentration in CO_2_-saturated water was calculated based on the temperature of the CO_2_ solution and atmospheric pressure. Water carbonation was conducted in a different room from the other experiments.

All *in vitro* experiments were carried out in atmospheric [CO_2_]; thus, some extent of atmospheric CO_2_ was dissolved in the control reactions. We considered applying a CO_2_-free atmosphere, but that could lead to increased release of CO_2_ from [CO_2_]_high_ reactions. The use of atmospheric CO_2_ partially limited CO_2_ loss from [CO_2_]_high_ reactions. Moreover, we maximally reduced the number of reactions prepared simultaneously to limit CO_2_ loss from [CO_2_]_high_ reactions.

### Plant growth

Arabidopsis WT Columbia-0 ecotype plants; mutant lines *ht1-2*^37^, *mpk4-2* (SALK_056245), *mpk3-1* (SALK_151594), and *mkk1 mkk2*; and a line expressing One-STrEP-tag-MPK4 were grown on soil in a GIR 96 growth chamber (Conviron, Winnipeg, Canada) at 22°C and 60–70% humidity under a 16-h light (100 μmol m^-2^ s^-1^)/8-h dark photoperiod.

### Preparation and treatment of epidermal peels

For 20 preparations, 300-350 rosette leaves (~45 g) excised from 3-week-old Arabidopsis plants were blended in 1,400 ml of demineralised water for 1.5 min. For 2 preparations from 5-week-old *mpk4-2* or *mkk1 mkk2* plants, 160-200 shoots were blended in 600 ml of demineralised water for 2 min. Epidermal peels were then collected on 100-μm Sefar Nitex mesh (Sefar AG, Heiden, Switzerland), washed three times with stomatal opening solution (20 mM MES-KOH (pH 5.7), 10 mM KCl, 50 μM CaCl_2_) and incubated in open tubes in 10 ml of stomatal opening solution for 3 h in a GIR 96 growth chamber. Under the indicated treatment, epidermal peels were retained on Sefar Nitex mesh and frozen in liquid nitrogen.

### Protein extraction from epidermal peels

Frozen epidermal peels (1.5 ml) were ground in a mortar upon the addition of 1.1 g of sucrose, 80 μl of 1.5 M Tris (pH 8.0), 80 μl of 20% SDS, 80 μl of β-mercaptoethanol and 4 ml of phenol equilibrated with 10 mM Tris-HCl (pH 8.0). The lysate was vortexed for 30 s, incubated for 3 min at RT and centrifuged (1 min, 500 x g, 4°C). The upper organic phase was transferred to 9 ml of isopropanol with 100 mM ammonium acetate. Proteins were precipitated at −20°C for 24 h and centrifuged (12,000 x g, 15 min, 4°C). The pellet was washed with 14 ml of methanol and then with 12 ml of ethanol and dried in air for 20 min at RT. Proteins were dissolved in 200 μl of Laemmli sample buffer with cOmplete EDTA-free Protease Inhibitor Cocktail (Roche, Mannheim, Germany) for 15 min at RT.

### One-STrEP-tag affinity purification

The coding sequences of the One-STrEP-tag fusion proteins under the control of the Arabidopsis *UBQ10* promoter and *NOS* terminator^35,36^, cloned in the binary vector pART27^38^, were stably expressed in Arabidopsis Col-0 plants following *Agrobacterium tumefaciens* (strain GV3101)-mediated transformation. Epidermal peels from 10 g of rosette leaves were ground in liquid nitrogen, resuspended in 3 ml of extraction buffer (100 mM Tris-HCl (pH 8.0), 200 mM NaCl, 100 mM NaF, 10 mM EDTA, 0.4% Triton X-100, 3 mM DTT, 3.2 mM Na_3_VO_4_, cOmplete EDTA-free Protease Inhibitor Cocktail (Roche, Mannheim, Germany)) and filtered on Sefar Nitex mesh (Sefar AG, Heiden, Switzerland). After centrifugation (13,000 x g, 4 min, 4°C), the supernatant was loaded onto Bio-Spin^®^ chromatography columns (Bio-Rad) containing 50 μl of Strep-Tactin Superflow high-capacity resin (IBA, Goettingen, Germany). After six washing steps (100 mM Tris-HCl (pH 8.0), 150 mM NaCl), proteins were eluted with 400 μl of 5 mM desthiobiotin (IBA, Goettingen, Germany) in washing solution, concentrated with Amicon Ultra 10K filters (Millipore, Billerica, MA, USA), aliquoted and stored at −80°C.

### High-resolution electrophoresis

Tris-glycine SDS-PAGE was carried out in a discontinuous buffer system with a 5% stacking gel (pH 6.8) and 9% resolving gel (pH 8.8). A total of 30-50 μg of total protein was loaded per lane of the gel (26 cm length, 14 cm width and 1 mm thickness). A step voltage reduction of 10 V every 10 min from 180 V to 140 V was applied during protein concentration in the stacking gel. In the resolving gel, electrophoresis was conducted at a constant current of 12 mA/gel (max. 180 V) for 16 h at room temperature.

### Immunoblotting

Denatured proteins separated on an 8-11.5% SDS-PAGE gel were transferred onto nitrocellulose membranes. The membranes were blocked for 60 min in 5% skimmed milk in TBST (20 mM Tris, 0.8% NaCl, 0.05% Tween-20) or 7% BSA in TBST and incubated at room temperature for 1 h with anti-MPK3, anti-MPK4, anti-MPK6 (1:500, Sigma-Aldrich, Steinheim, Germany), anti-thiophosphate ester (anti-TE, ab92570, 1:5,000, Abcam, Cambridge, UK), anti-phospho-MBP (13-104, 1:200, Merck) or phospho-p44/42 MAPK (Erk1/2) (Thr202/Tyr204) antibody (anti-phospho-TEY, #9101, 1:200, Cell Signaling Technology, Danvers, MA, USA). Membranes were washed 3 times for 5 min with TBST and incubated for 1 h with the appropriate secondary antibody – goat anti-rabbit (1:20,000, Agrisera, Vännäs, Sweden) or goat anti-mouse (1:160,000 Thermo Scientific, Rockford, IL, USA). Detection was performed with ECL (Thermo Scientific, Rockford, IL, USA) according to the manufacturer’s instructions.

### *In vitro* MPK activity measurement

Due to the tendency of active MPK4 to aggregate^11^, MPK4 was diluted to working concentration in CO_2_-free water containing 2.5 mM DTT and, when indicated, 1 mg/ml BSA (0.4 mg ml^-1^ in *in vitro* reaction) as an antiaggregatory factor^39^. One-STrEP-tagged kinases purified from Arabidopsis epidermal peels or 0.2-1 μg of kinases overexpressed in bacteria was incubated (25 min at 30°C or as indicated) with 2.5 μg of MBP (Millipore, Temecula, CA, USA) or 0.5 μg of another substrate protein, as indicated, in buffer containing 40 mM MOPS (pH 7.0 or as indicated), 0.5 mM EGTA, 1 mM DTT, 20 mM MgCl_2_, 200 μM ATP and Protease and Phosphatase Inhibitor Tablets, EDTA Free (Thermo Scientific, Rockford, IL, USA). When protein thiophosphorylation was detected by immunoblotting with anti-TE^40^, reactions were performed in buffer containing 40 mM MOPS (pH 7.0 or as indicated), 0.5 mM EGTA, 1 mM DTT, 20 mM MgCl_2_, 1 ATP-γ-S (adenosine-5’-O-(3-thiotriphosphate), BIOLOG Life Science Institute, Bremen, Germany) and Protease and Phosphatase Inhibitor Tablets, EDTA Free (Thermo Scientific, Rockford, IL, USA). After thiophosphorylation, 2.5 mM *p*-nitrobenzyl mesylate (Abcam, Cambridge, UK) was added, and the samples were further incubated for 25 min at room temperature. Then, proteins were separated by SDS-PAGE and subjected to immunoblotting with anti-TE, anti-phospho-MBP or anti-phospho-TEY.

### CO_2_ binding assay

Four micrograms of MPKs was incubated for 10 min at 24°C in 100 μl of binding reaction containing 100 mM MOPS (pH 6.4-7.0 as indicated), 5 mM EGTA, 20 mM MgCl_2_, 1 mM DTT, 200 μM ATP (optional), cOmplete EDTA-free Protease Inhibitor Cocktail (Roche, Mannheim, Germany) and the indicated concentration of [^14^C] KHCO_3_ (50-60 mCi mmol ^-1^). Then, the samples were vortexed for 30 s and loaded onto 2 ml of Sephadex G-25 coarse (Pharmacia) in Bio-Spin^®^ chromatography columns and washed with 240 μl of washing buffer containing 20 mM MES (pH 6.4-7.0 according to the pH of the binding reaction), 2% BSA, 20 mM MgCl_2_, 2 mM DTT, and 50 mM NaCl. Then, proteins were eluted using washing buffer. Two 100-μl fractions were collected in 50 μl of 1.5 M Tris-HCl (pH 8.0) and 1 ml of Ultima Gold LLT scintillation cocktail (Perkin Elmer). Radioactivity was measured using a liquid scintillation analyser (Packard).

### Barley mesophyll protoplast transformation

Twenty 6- to 7-day-old barley leaves were sliced crosswise to obtain scraps with minimal thickness. Sliced material was incubated in 60 ml of enzyme solution (0.615 M mannitol, 1.5% Cellulase Onozuka R10 (Serva, Heidelberg, Germany), 0.3% Macerozyme R10 from *Rhizopus sp*. (Serva, Heidelberg, Germany), 1% BSA, 10 mM MES; pH 5.7) for 3 h at 28°C. After slow cooling (20 min at 4°C), the suspension was gently swirled to facilitate protoplast release and filtered through 100-μm Sefar Nitex mesh. Subsequent stages were carried out on ice or at 4°C. Protoplasts were centrifuged at 250 x g for 4 min and washed twice with 0.615 M mannitol. Protoplasts (2×10^5^) resuspended in 0.615 M mannitol were added to 40 μg of individual plasmids in 0.615 M mannitol in a final volume of 110 μl. Electroporation was carried out using Gene Pulser Xcell (Bio-Rad) in 4-mm electroporation cuvettes (Bio-Rad) with the following setting: a single pulse at 150 V with an 8-ms pulse duration. Immediately, 1 ml of ice-cold 0.615 M mannitol was added, and protoplasts were transferred to 2-ml tubes. The protoplasts were allowed to sediment for 20 min at room temperature before resuspension in incubation solution (0.615 M mannitol (pH 5.9), 10 mM CaCl_2_, 1 mM MgSO_4_, 1 mM KNO_3_, 100 μM KH_2_PO_4_, 10 μM KI, 1 μM CuSO_4_). Protein localization was documented after overnight incubation at 21 °C.

### Arabidopsis mesophyll protoplast transformation

The epidermis from the underside of 6-7 rosette leaves (from 4-5-week-old plants) was peeled away using Scotch Magic Tape 3M adhered to both sides of the leaf^41^. Leaves were incubated in Petri dishes with 10 ml of enzyme solution (1.2% Cellulase Onozuka R10, 0.4% Macerozyme R10, 0.4 M mannitol, 20 mM KCl, 20 mM MES; pH 5.7) for 90 min at room temperature with gentle rotation (20 rpm on a platform shaker). Protoplasts were then diluted (1:1) with ice-cold W5 solution (154 mM NaCl, 125 mM CaCl2, 5 mM KCl, 2 mM MES; pH 5.7), centrifuged (3 min, 100 x g, 4°C) and washed twice with 30 ml of ice-cold W5 solution. After the last wash, protoplasts were allowed to sediment on ice for 30 min and resuspended in MMg solution (0.4 M mannitol, 15 mM MgCl_2_, 4 mM MES; pH 5.7) to obtain a concentration of 2×10^4^ cells ml^-1^. For transfection^42^, 100 μl of protoplasts were transferred to wells of U96 Microwell plates (Thermo Scientific Nunc), mixed (700 rpm) with 5 μg of plasmids (in 10 μl) and 110 μl of PEG solution (40% PEG 4,000, 200 mM mannitol, 100 mM CaCl_2_), and incubated for 5 min (with 10 s of mixing at intervals of 50 s) at room temperature. Then, 200 μl of W5 solution was added and mixed (800 rpm for 10 s). Protoplasts were centrifuged at 100 x g for 1 min and washed 4 times with W5 solution (200 μl of solution from the wells was removed, and 200 μl of fresh W5 was added, mixed for 10 s at 800 rpm and centrifuged at 100 x g for 1 min). Protoplasts were incubated in a growing chamber for 12-16 h.

### Arabidopsis guard cell protoplast transformation

Epidermises from the undersides of 12 rosette leaves (from 4- to 5-week-old plants) were incubated in Petri dishes with 10 ml of enzyme solution (1.8% Cellulase Onozuka R10, 0.8% Macerozyme R10, 0.4 M mannitol, 20 mM KCl, 20 mM MES; pH 5.7) at room temperature with gentle rotation (20 rpm on a platform shaker) until all mesophyll and most pavement cells peeled off. Guard cells bound to the Scotch Magic Tape were transferred to a fresh portion of enzyme solution and digested for 30-45 min. Protoplasts were then centrifuged (5 min, 450 x g) and washed with 30 ml of MMg. Finally, MMg was added to obtain 1×10^5^ cells ml^-1^. All subsequent steps were carried out as described for mesophyll protoplasts, with modified centrifugation steps (300 x g, 2 min). Guard cell protoplasts were incubated in modified W1 solution (0.5 M mannitol, 15 mM KCl, 50 μM CaCl_2_ and 10 mM MES-Tris; pH 6.15) in a growing chamber for 12-16 h.

### Protein localization

Arabidopsis guard cell or mesophyll protoplasts and barley mesophyll protoplasts were transfected with plasmids encoding proteins fused to EYFP (pSAT4A-EYFP-N1 vector^43^). After transfection, protoplasts were transferred to black 96-well black glass-bottom plates (SensoPlate, Greiner Bio-One) and incubated overnight in a growth chamber. Protein localization was documented with a Nikon A1Rsi confocal system with the following settings: dichroic mirror, 457/514; A1-DU4 4 detector unit; filter, 540/30. An argon ion laser (514 nm, laser power: 0.8) was used for excitation of EYFP.

### Plasmid construction

All plasmids used were modified such that *Sfi*I restriction sites (arranged as in the pUNI51 vector, GenBank accession AY260846) were placed into their polylinkers. pUNI51 clones containing coding sequences of MPK2 (U10062), MPK4 (U09192), MPK6 (U15193), MPK12 (U82548) and MPK20 (U13519) were obtained from the Arabidopsis Biological Resource Center (Columbus, OH, USA). Other coding sequences were amplified from Arabidopsis or barley cDNAs and cloned in pUNI51. Plasmids encoding GST-fusion proteins were constructed in pGEX-6P-1 (MPK4, MPK4 mutants) or by loxP/Cre-based recombination of pHB2-GST and pUNI51 plasmids^44^ (other MPKs).

### Statistical analysis

The presented statistically significant differences in results from at least three experiments (means ± standard deviations) were based on one-way or factorial ANOVA, followed by Tukey’s post hoc comparison.

ImageJ software^45^ was employed for densitometric analysis of immunoblotting bands. Kinase activities were calculated in terms of protein amounts. For data normalization, the sum of all kinase activity measurements throughout the experiment was taken as 1. Then, for clarity, the value of the control was taken as 1.

## Acknowledgements

The authors thank Mirosława Dabert and Wiesława Jarmuszkiewicz for logistic and mentoring support, Adam Augustyniak and Michał Kopa for technical assistance and Dawid Bielewicz, Tomasz Bieluszewski, Koh Iba and Yuelin Zhang for providing seeds. This work was supported by the National Science Center, Poland (UMO-2011/01/D/NZ3/02068 to ŁG and UMO-2015/19/D/NZ3/00479 to HG).

## Author Contributions

HG and ŁG: Conceptualization, funding acquisition, experiment execution, data analysis, and writing.

## Competing interests

The authors declare no competing interests.

---

**Supplementary Fig. 1.**
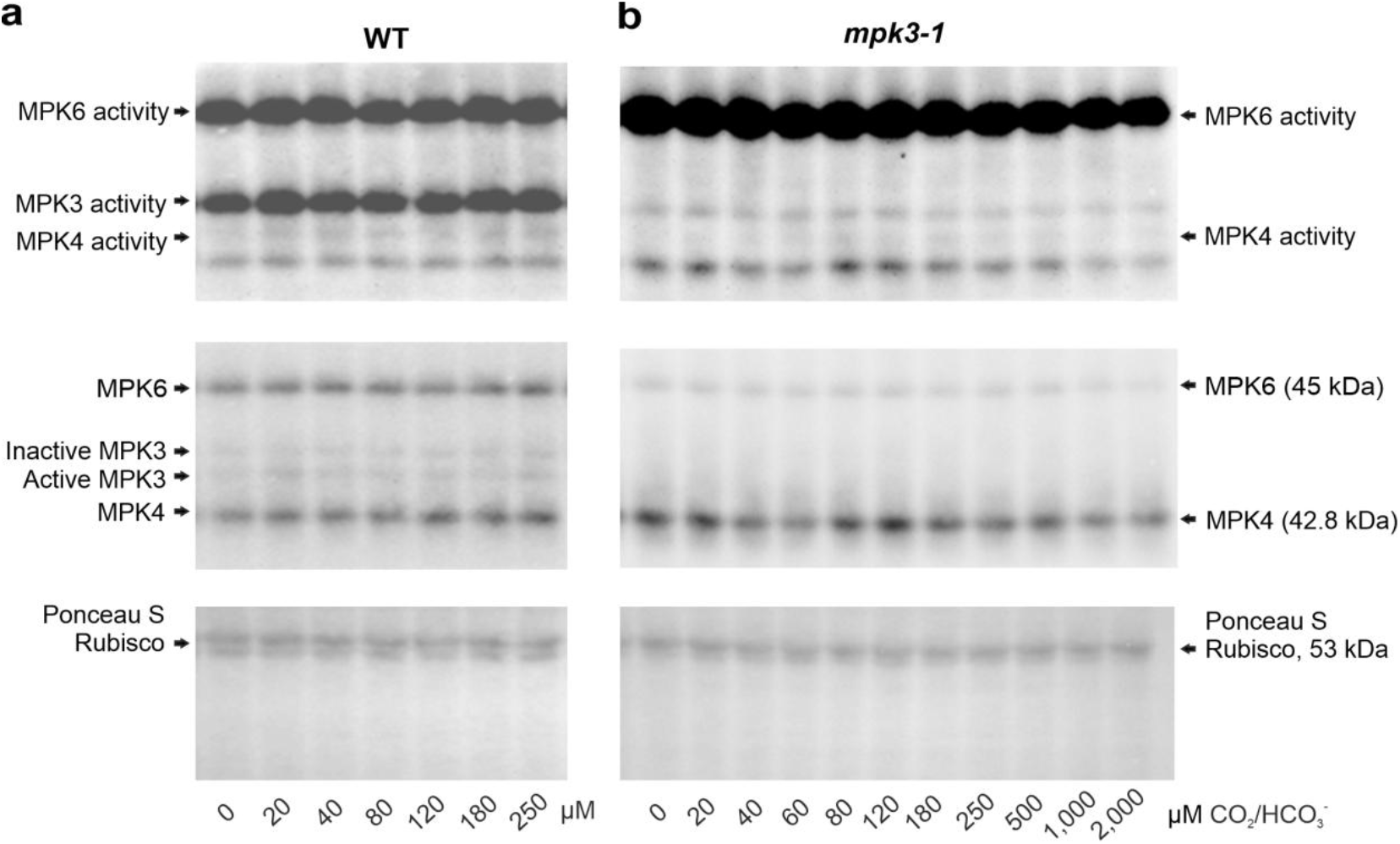
Additional experiments showing MPK4 activation by CO_2_ in epidermal peels. **a**, Similar to dissolved CO_2_, an increase in [HCO_3_^-^] induces MPK4 activity. **b**, A study on MPK activity in epidermal peels of *mpk3-1* showed that strong immunoblotting signals from MPK3 did not influence the measurement of MPK4 activity. Before administration of the indicated [HCO_3_^-^], epidermal peels were incubated at pH 5.7 in open tubes ensuring stabilization of [CO_2_], which may fluctuate due to CO_2_ consumption and production by epidermal peels. Then, the indicated [HCO_3_^-^] was added for 15 min. Active MPKs were immunodetected with anti-phospho-TEY following SDS-PAGE. Protein loading was assessed by Ponceau S staining and immunoblotting with anti-MPK3, anti-MPK4 and anti-MPK6 antibodies. Data from two biological replicates.

**Supplementary Fig. 2.**
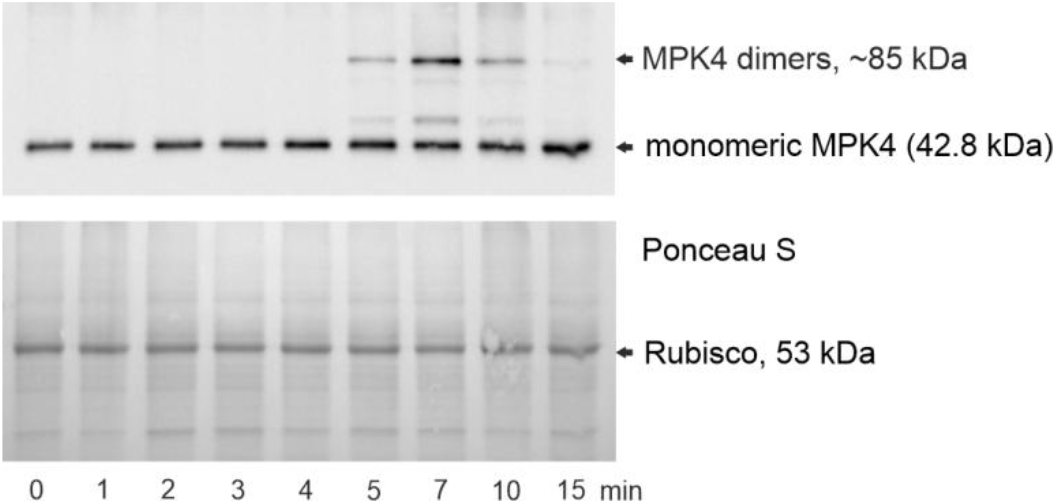
An additional MPK4 band was recognized in response to exposure to 180 μM HCO_3_^-^. Due to the double molecular mass compared to monomeric MPK4 and known MPK4 susceptibility to multimerization, we expect that the most likely modification of ~85-kDa MPK4 is covalent dimerization. Protein bands of the same molecular weight, emerging time points and intensity profile were detected using anti-TEY and anti-MPK4 antibodies (Fig. 1c), indicating that dimeric MPK4 is ~85-kDa active MPK. The amounts of proteins were determined by staining with Ponceau S and immunoblotting with anti-MPK4. Representative results from three experiments are presented.

**Supplementary Fig. 3.**
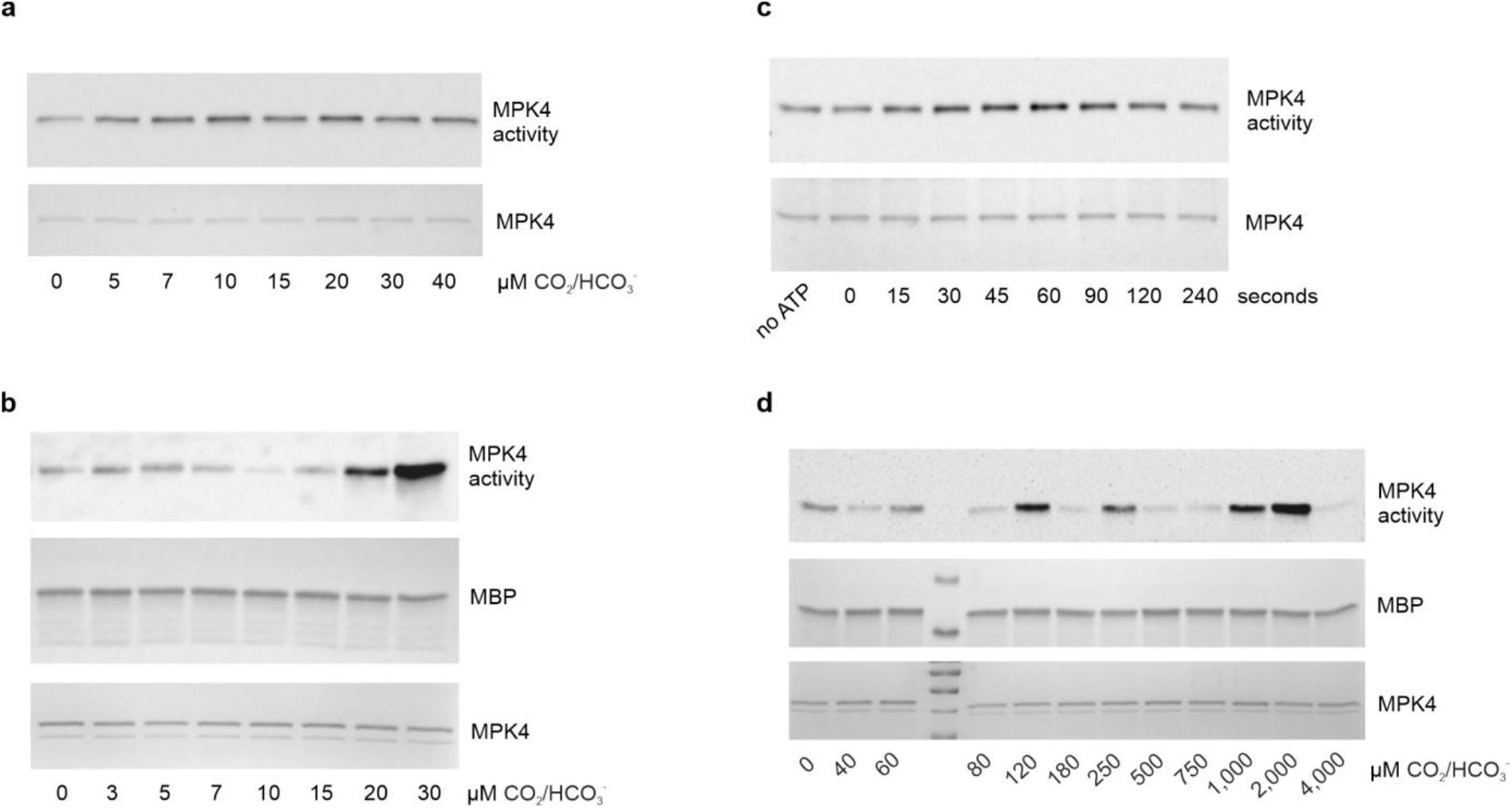
Investigation of *in vitro* MPK4 activation by CO_2_/HCO_3_^-^ (at pH 7.0). **a**, **b**, Similar to dissolved CO_2_, KHCO_3_ regulates MPK4 activity. **c**, MPK4 activation in response to [CO_2_]_high_ occurs in just a few seconds. **d**, Very high CO_2_/HCO_3_^-^ concentrations can positively or negatively regulate MPK4 activity. The lack of MPK4 activation in response to millimolar CO_2_/HCO_3_^-^ concentration is consistent with a previous report^2^. In **a**, **c**, MPK4 activity is shown by immunoblotting with anti-phospho-TEY and MPK4 loading by staining with Ponceau S. In **b**, **d**, MPK4 activity was determined by MBP *in vitro* thiophosphorylation and detected by immunoblotting with anti-TE. MPK4 was stained with CBB, and MBP was stained with Ponceau S. Experiments were carried out using MPK4 purified from bacteria and dephosphorylated by FastAP phosphatase; GST-MPK4 (**c**), tag-free MPK4 (**a-b, d**). Representative results from three independent experiments are presented.

**Supplementary Fig. 4.**
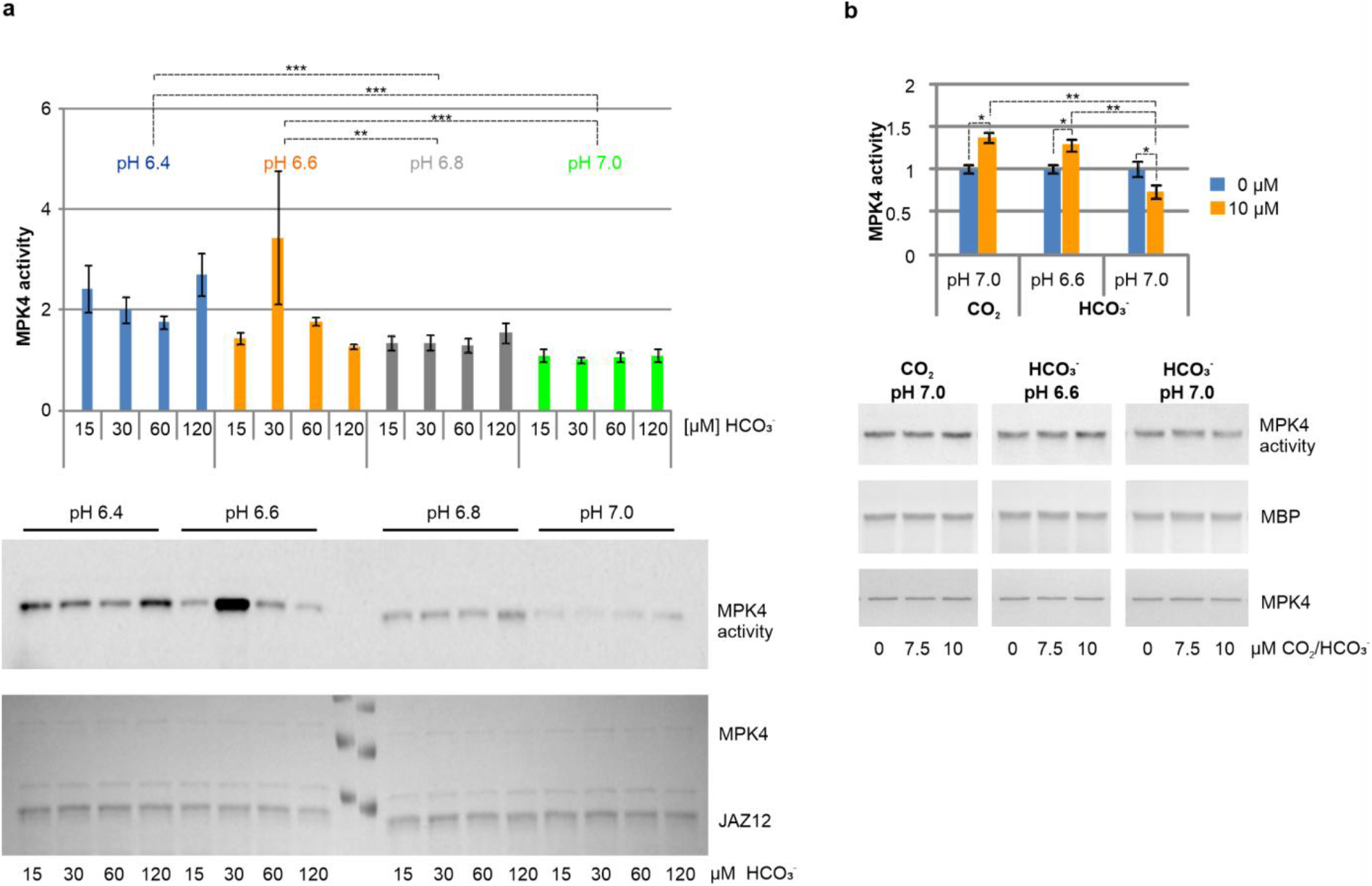
Increase in [CO_2_] enhances MPK4 kinase activity, and increase in [HCO_3_^-^] reduces MPK4 kinase activity. **a**, Elevation of [HCO_3_^-^] at constant [CO_2_] triggers MPK4 inactivation. *In vitro* thiophosphorylation reactions with different [CO_2_/HCO_3_^-^] in the pH series were allowed exchange with ambient air for 30 min, leading to equalization of [CO_2_] in all samples so that they differed in only [HCO_3_^-^]. Then, MPK4 and ATPγS were added, and *in vitro* thiophosphorylation reactions were carried out for only 2 min. High pH and concomitant high [HCO_3_^-^] led to low MPK4 activity. JAZ12 was thiophosphorylated by dephosphorylated GST-MPK4, and its activity was detected using immunoblotting with anti-TE. JAZ12 and MPK4 bands were stained by Ponceau S. **b**, MPK4 is activated by 7.5-10 μM CO_2_ at pH 7.0 but inactivated by 7.5-10 μM HCO_3_^-^. Lowering the pH to 6.6 (increase in free [CO_2_] and decrease in [HCO_3_^-^]) leads to reversal of the HCO_3_^-^-induced MPK4 activity profile to that triggered by dissolved CO_2_. MBP was used as a substrate of tag-free MPK4, and MPK4 activity was detected by immunoblotting with anti-phospho-MBP. MBP loading was visualized by Ponceau S, and MPK4 loading was visualized by CBB. Representative data from three experiments. Mean ±SD; *, ** and *** indicate p<0.05, p<0.01 and p<0.001, respectively.

**Supplementary Fig. 5.**
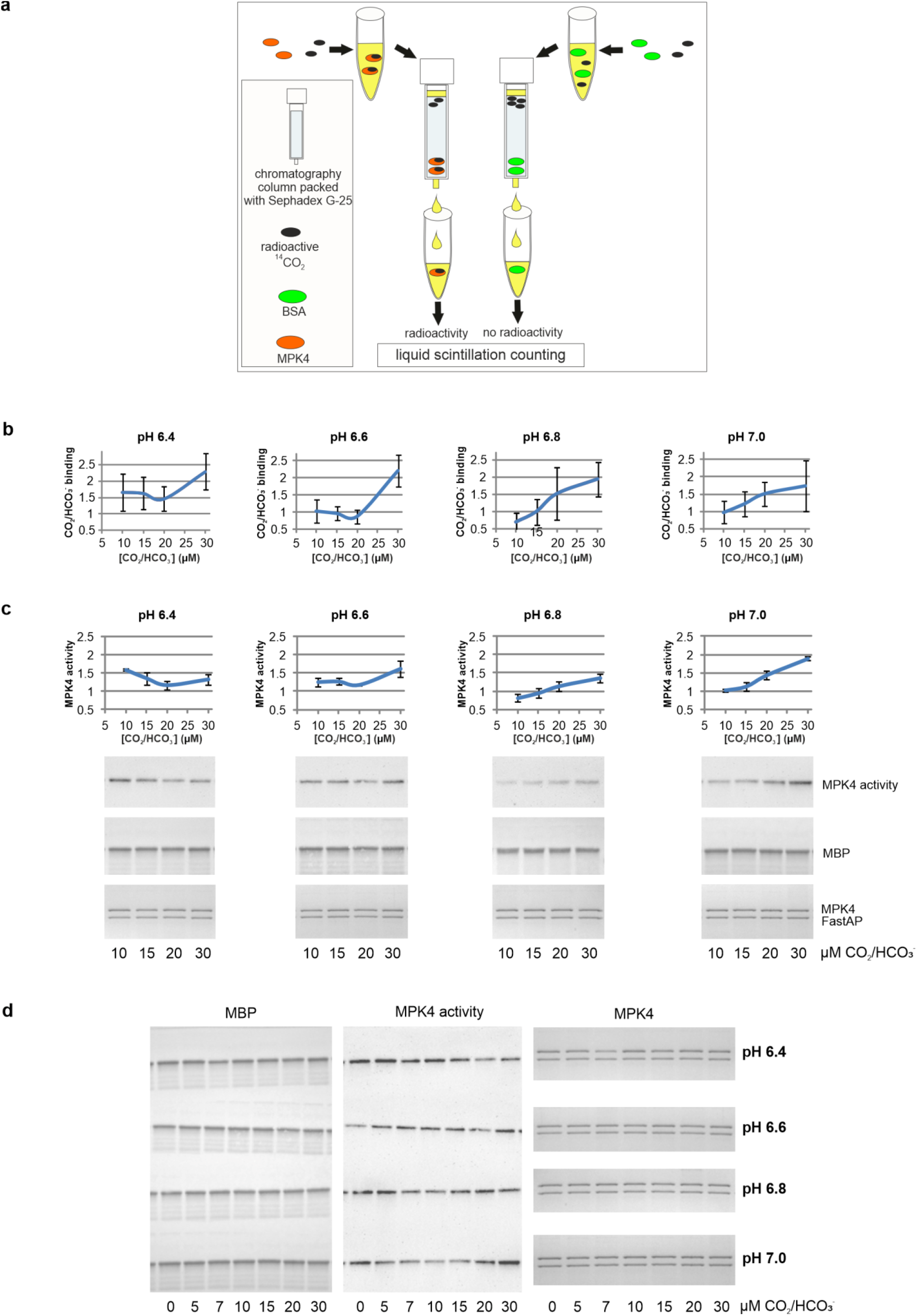
Increase in MPK4 activity is correlated with enhanced CO_2_ binding. **a**, Scheme of gel filtration-based CO_2_ binding assay. **b**, Graphs of CO_2_ binding at 10-30 μM CO_2_/HCO_3_^-^ in individual pH series from the graph shown in Fig. 3b; data normalization was based on values of no-protein controls from each experiment; mean ±SD, n=3 experiments. **c**, MPK4 activity under the conditions applied for the CO_2_ binding assay shown in **b**. Dephosphorylated MBP was used as a substrate of dephosphorylated tag-free MPK4; FastAP-fast alkaline phosphatase. The intensity of MBP phosphorylation was detected by immunoblotting with anti-phospho-MBP. The amount of MBP was determined by Ponceau S, and the amount of MPK4 was determined by CBB. Mean ± SD, n=3 experiments. **d**, Example original images of immunoblotting and protein staining used to calculate data for graphs presented in **c**.

**Supplementary Fig. 6.**
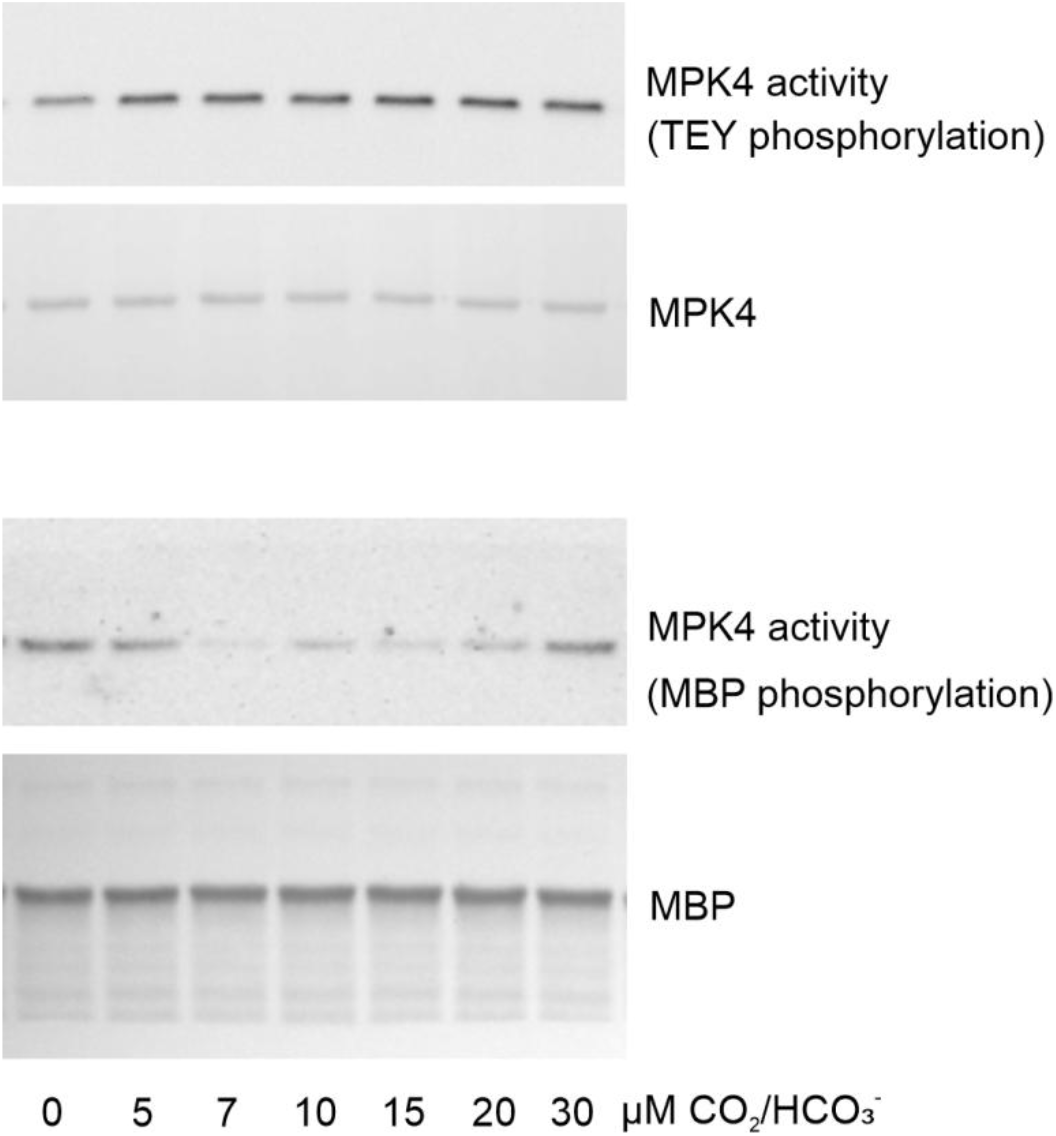
[CO_2_]_hig_h-induced TEY phosphorylation is not inhibited by HCO_3_^-^ at pH ≥7, unlike the decrease in MPK4 activity, defined as substrate phosphorylation intensity. TEY and MBP phosphorylation is shown by immunoblotting with anti-phospho-TEY and anti-phospho-MBP, respectively. Both analyses were carried out from one set of *in vitro* phosphorylation reactions. The amounts of dephosphorylated MBP and dephosphorylated MPK4 on the nitrocellulose membrane were specified by Ponceau S staining. Representative images from three experiments.

**Supplementary Fig. 7.**
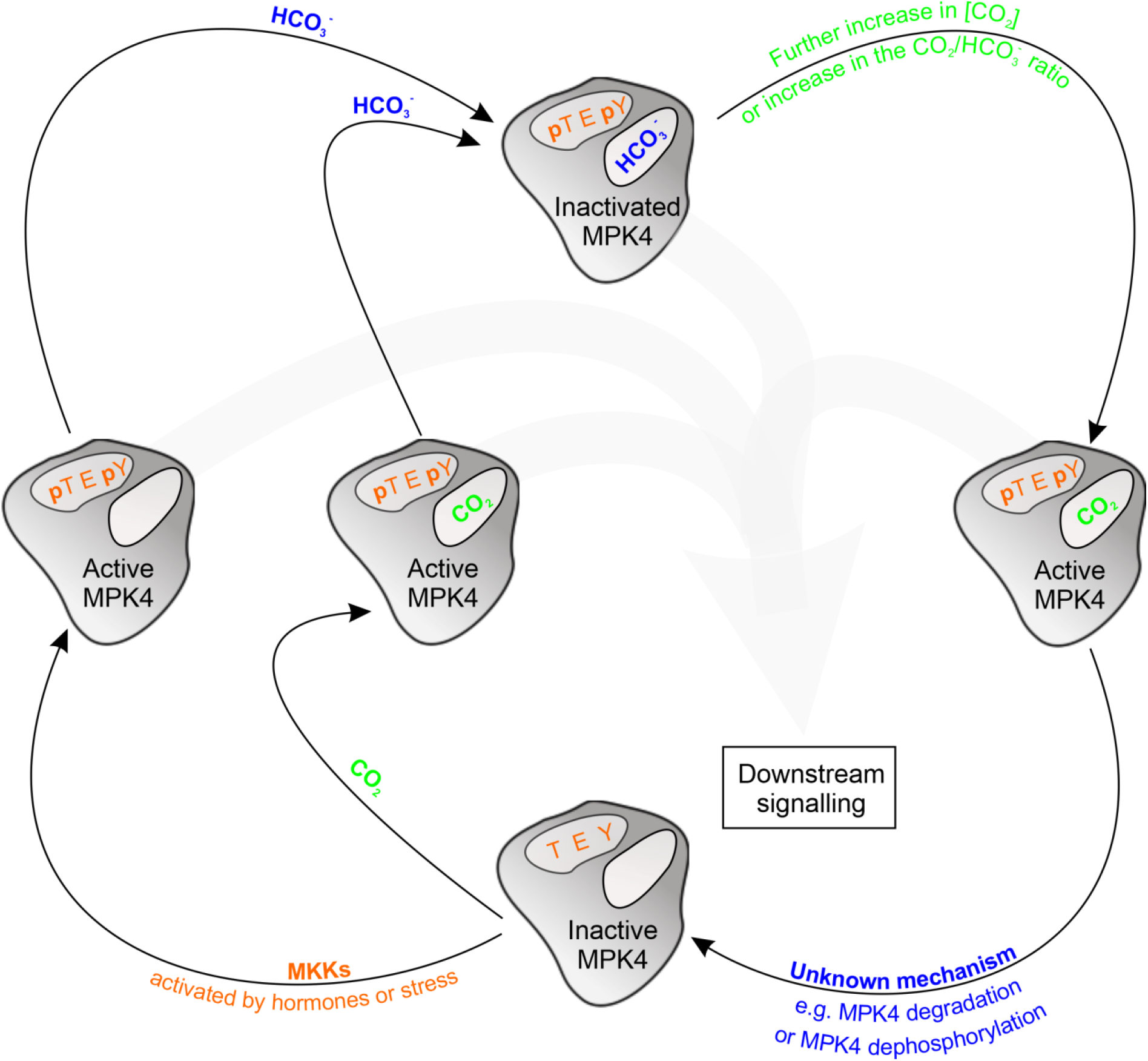
Proposed working model of the MPK4 response to [CO_2_/HCO_3_^-^]high.

**Supplementary Fig. 8.**
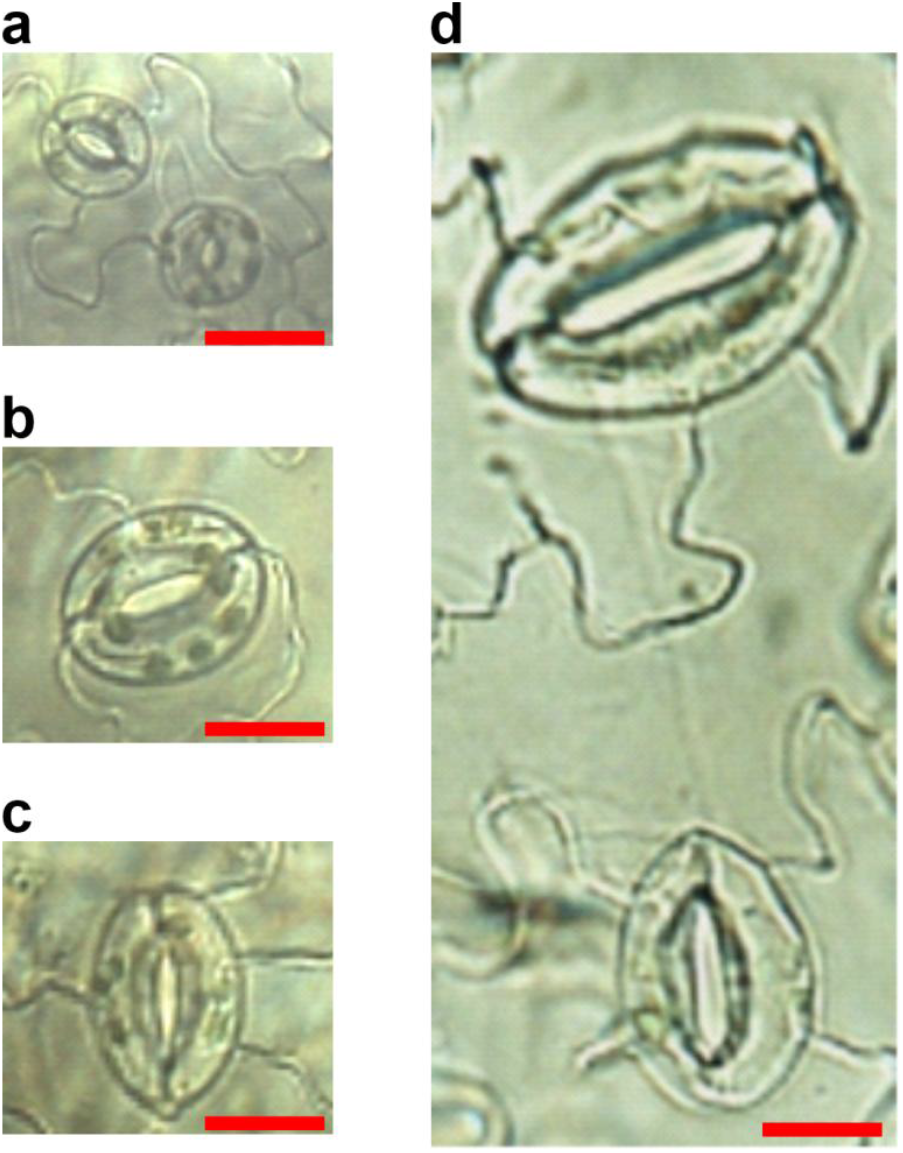
MPK4 influences stomatal development. **a**, Stomata of WT Arabidopsis. **b-d**, Enlarged and elongated stomata in *mpk4-2* leaves. Scale bars 20 μm. As reported for stomata of an *N. tabacum* line with silenced *NtMPK4*^1^, *mpk4-2* stomata display a much wider range of length than WT stomata.

**Supplementary Fig. 9.**
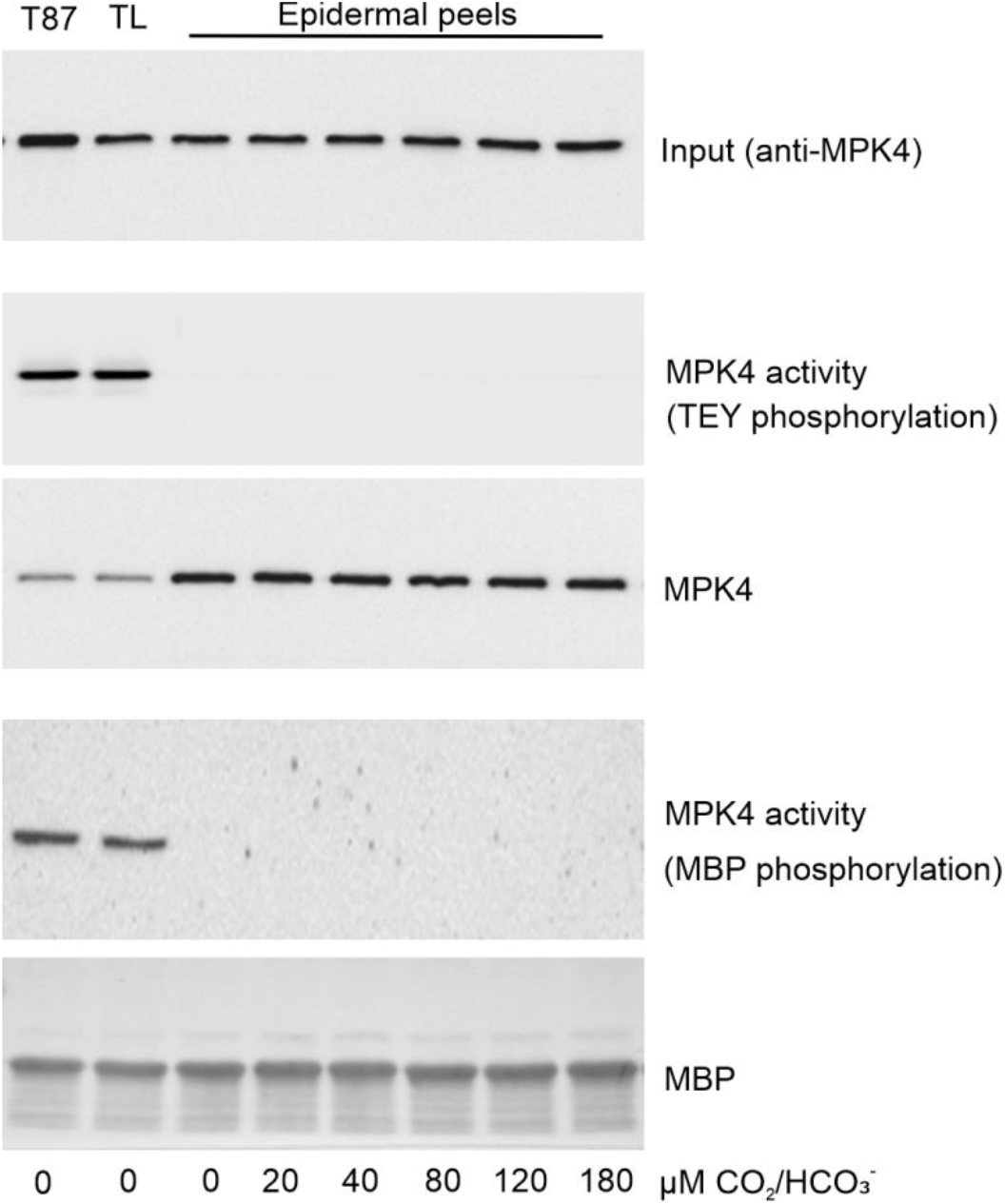
Inactive One-STrEP-Tag-MPK4 was specifically purified from Arabidopsis epidermal peels under native conditions. Anti-phospho-TEY antibody and MBP *in vitro* phosphorylation experiments failed to detect the activity of guard cell One-STrEP-Tag-MPK4 in contrast to One-STrEP-Tag-MPK4 from Arabidopsis total leaf (TL) extracts or T87 cultured cells. We used a powerful method for specific purification of One-STrEP-tagged plant proteins under native conditions within several minutes^35,36^. In contrast to high-yield One-STrEP-Tag-MPK4 purification from epidermal peels, we were not able to detect One-STrEP-Tag-MPK4 activity by *in vitro* MBP phosphorylation. Based on the membrane-associated localization of barley MPK4 (Supplementary Fig. 10b), we hypothesize that MPK4 activatable by [CO_2_]_hig_h is connected to the cell membrane. In addition, the use of phenol-SDS extraction (Fig. 1, Supplementary Fig. 1), which increases membrane protein solubilization and decreases protein interactions, underlies the successful detection of MPK4 activity.

**Supplementary Fig. 10.**
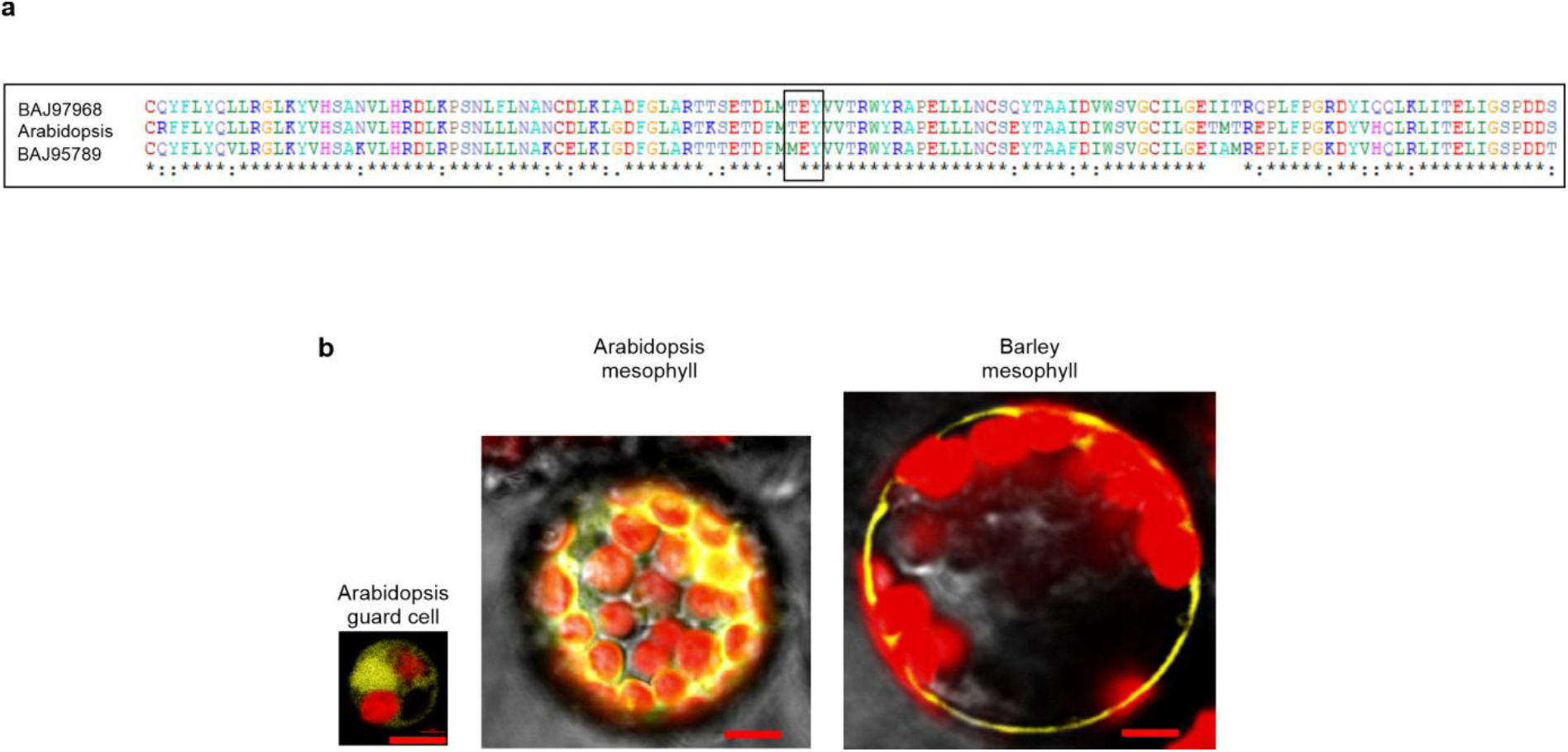
**a**, Alignment (using ClustalX 2.1) of amino acid sequences of Arabidopsis MPK4 and its two barley homologues. The response of barley stomata to darkness is the quickest among the studied species^34^. It may be speculated that this results from the presence of two specialized MPK4 homologues in barley guard cells. The protein encoded by the BAJ95789 locus shares lower identity (82%) with Arabidopsis MPK4 than the MPK encoded by BAJ97968 (83%). Moreover, the polypeptide encoded by BAJ95789 does not contain the TEY motif (in a black frame); therefore, the barley MPK encoded in the BAJ97968 locus was included in the comparative analysis of [CO_2_]_high_-induced MPK activity in Fig. 4a, and the expression in barley protoplasts is presented in **b**. **b**, Barley MPK4-YFP in barley mesophyll protoplasts is localized in the proximity of the cell membrane, in contrast to Arabidopsis MPK4-YFP, which was predominantly dispersed in the cytoplasm and nucleus in both the Arabidopsis mesophyll and Arabidopsis guard cell protoplasts. Bar, 2.5 μm.

## Notes

### Competing Interest Statement

The authors have declared no competing interest.

## References

1. Marten, H. et al. Silencing of *Nt* MPK4 impairs CO_2_-induced stomatal closure, activation of anion channels and cytosolic Ca^2+^ signals in *Nicotiana tabacum* guard cells. Plant J. 55, 698–708 (2008).

2. Tõldsepp, K. et al. Mitogen-activated protein kinases MPK4 and MPK12 are key components mediating CO_2_-induced stomatal movements. Plant J. 96, 1018–1035 (2018).

3. Hõrak, H. et al. A Dominant Mutation in the HT1 Kinase Uncovers Roles of MAP Kinases and GHR1 in CO_2_-Induced Stomatal Closure. Plant Cell 28, 2493–2509 (2016).

4. Petersen, M. et al. Arabidopsis MAP Kinase 4 Negatively Regulates Systemic Acquired Resistance. Cell 103, 1111–1120 (2000).

5. Hsu, P.-K. et al. Abscisic acid-independent stomatal CO_2_ signal transduction pathway and convergence of CO_2_ and ABA signaling downstream of OST1 kinase. Proc. Natl. Acad. Sci. 115, E9971–E9980 (2018).

6. He, J. et al. The BIG protein distinguishes the process of CO_2_-induced stomatal closure from the inhibition of stomatal opening by CO_2_. New Phytol. 218, 232–241 (2018).

7. Tian, W. et al. A molecular pathway for CO_2_ response in Arabidopsis guard cells. Nat. Commun. 6, 6057 (2015).

8. Hiyama, A. et al. Blue light and CO_2_ signals converge to regulate light-induced stomatal opening. Nat. Commun. 8, 1284 (2017).

9. Jakobson, L. et al. Natural Variation in Arabidopsis Cvi-0 Accession Reveals an Important Role of MPK12 in Guard Cell CO_2_ Signaling. PLOS Biol. 14, e2000322 (2016).

10. Raschke, K. Saturation Kinetics of the Velocity of Stomatal Closing in Response to CO_2_. Plant Physiol. 49, 229–34 (1972).

11. Zhang, T., Zhu, M., Song, W., Harmon, A. C. & Chen, S. Oxidation and phosphorylation of MAP kinase 4 cause protein aggregation. Biochim. Biophys. Acta - Proteins Proteomics 1854, 156–165 (2015).

12. Berghuijs, H. N. C. et al. Localization of (photo)respiration and CO_2_ re-assimilation in tomato leaves investigated with a reaction-diffusion model. PLoS One 12, e0183746 (2017).

13. Hu, H. et al. Carbonic anhydrases are upstream regulators of CO_2_-controlled stomatal movements in guard cells. Nat. Cell Biol. 12, 87–93 (2010).

14. DiMario, R. J. et al. The Cytoplasmic Carbonic Anhydrases βCA2 and βCA4 Are Required for Optimal Plant Growth at Low CO_2_. Plant Physiol. 171, 280–93 (2016).

15. Kosetsu, K. et al. The MAP kinase MPK4 is required for cytokinesis in *Arabidopsis thaliana*. Plant Cell 22, 3778–90 (2010).

16. Gawroński, P. et al. Mitogen-activated protein kinase 4 is a salicylic acid-independent regulator of growth but not of photosynthesis in Arabidopsis. Mol. Plant 7, 1151–66 (2014).

17. Imai, Y. et al. Angiotensin-converting enzyme 2 protects from severe acute lung failure. Nature 436, 112–6 (2005).

18. Chen, I.-Y. et al. Upregulation of the chemokine (C-C motif) ligand 2 via a severe acute respiratory syndrome coronavirus spike-ACE2 signaling pathway. J. Virol. 84, 7703–12 (2010).

19. Meng, Y. et al. Angiotensin-converting enzyme 2/angiotensin-(1-7)/Mas axis protects against lung fibrosis by inhibiting the MAPK/NF-κB pathway. Am. J. Respir. Cell Mol. Biol. 50, 723–36 (2014).

20. Li, Y. et al. Angiotensin-converting enzyme 2 prevents lipopolysaccharide-induced rat acute lung injury via suppressing the ERK1/2 and NF-κB signaling pathways. Sci. Rep. 6, 27911 (2016).

21. Li, Y. et al. Angiotensin-converting enzyme inhibition attenuates lipopolysaccharide-induced lung injury by regulating the balance between angiotensin-converting enzyme and angiotensin-converting enzyme 2 and inhibiting mitogen-activated protein kinase activation. Shock 43, 395–404 (2015).

22. Hung, Y.-H. et al. Alternative Roles of STAT3 and MAPK Signaling Pathways in the MMPs Activation and Progression of Lung Injury Induced by Cigarette Smoke Exposure in ACE2 Knockout Mice. Int. J. Biol. Sci. 12, 454–65 (2016).

23. Lin, C.-I. et al. Instillation of particulate matter 2.5 induced acute lung injury and attenuated the injury recovery in ACE2 knockout mice. Int. J. Biol. Sci. 14, 253–265 (2018).

24. Zhou, P. et al. A pneumonia outbreak associated with a new coronavirus of probable bat origin. Nature 579, 270–273 (2020).

25. Lan, J. et al. Structure of the SARS-CoV-2 spike receptor-binding domain bound to the ACE2 receptor. Nature (2020). doi:10.1038/s41586-020-2180-5

26. Schuh, K. & Pahl, A. Inhibition of the MAP Kinase ERK Protects From Lipopolysaccharide-Induced Lung Injury. Biochem. Pharmacol. 77, (2009).

27. Tang, S.-E. et al. Pre-Treatment with Ten-Minute Carbon Dioxide Inhalation Prevents Lipopolysaccharide-Induced Lung Injury in Mice via Down-Regulation of Toll-Like Receptor 4 Expression. Int. J. Mol. Sci. 20, (2019).

28. Schmetterer L, Lexer F, Findl O, Graselli U, Eichler HG, W. M. The Effect of Inhalation of Different Mixtures of O_2_ and CO_2_ on Ocular Fundus Pulsations. Exp. Eye Res. 63, 351–355 (1996).

29. Ohlraun, S. et al. CARbon DIoxide for the treatment of Febrile seizures: rationale, feasibility, and design of the CARDIF-study. J. Transl. Med. 11, 157 (2013).

30. Szollosi, I. et al. Effect of CO_2_ Inhalation on Central Sleep Apnea and Arousals From Sleep. Respiration. 71, (2004).

31. Baddeley, H. et al. Gas exchange parameters in radiotherapy patients during breathing of 2%, 3.5% and 5% carbogen gas mixtures. Br. J. Radiol. 73, 1100–1104 (2000).

32. Bradley, S. M., Simsic, J. M. & Atz, A. M. Hemodynamic effects of inspired carbon dioxide after the Norwood procedure. Ann. Thorac. Surg. 72, 2084–2088 (2001).

33. Slater, E. C., Rosing, J. & Mol, A. The phosphorylation potential generated by respiring mitochondria. Biochim. Biophys. Acta - Bioenerg. 292, 534–553 (1973).

34. Elliott-Kingston, C. et al. Does Size Matter? Atmospheric CO_2_ May Be a Stronger Driver of Stomatal Closing Rate Than Stomatal Size in Taxa That Diversified under Low CO_2_. Front. Plant Sci. 7, 1253 (2016).

35. Ludwików, A. et al. Arabidopsis protein phosphatase 2C ABI1 interacts with type I ACC synthases and is involved in the regulation of ozone-induced ethylene biosynthesis. Mol. Plant 7, 960–976 (2014).

36. Bieluszewski, T. et al. AtEAF1 is a potential platform protein for Arabidopsis NuA4 acetyltransferase complex. BMC Plant Biol. 15, 75 (2015).

37. Hashimoto, M. et al. Arabidopsis HT1 kinase controls stomatal movements in response to CO_2_. Nat. Cell Biol. 8, 391–7 (2006).

38. Gleave, A. A Versatile Binary Vector System With a T-DNA Organisational Structure Conducive to Efficient Integration of Cloned DNA Into the Plant Genome. Plant Mol. Biol. 20, 1203–1207 (1992).

39. Finn, T. E., Nunez, A. C., Sunde, M. & Easterbrook-Smith, S. B. Serum Albumin Prevents Protein Aggregation and Amyloid Formation and Retains Chaperone-like Activity in the Presence of Physiological Ligands. J. Biol. Chem. 287, 21530–21540 (2012).

40. Allen, J. J. et al. A semisynthetic epitope for kinase substrates. Nat. Methods 4, 511–6 (2007).

41. Wu, F.-H. et al. Tape-Arabidopsis Sandwich - a simpler Arabidopsis protoplast isolation method. Plant Methods 5, 16 (2009).

42. Fujikawa, Y. & Kato, N. Split luciferase complementation assay to study protein-protein interactions in Arabidopsis protoplasts. Plant J. 52, 185–95 (2007).

43. Tzfira, T. et al. pSAT vectors: a modular series of plasmids for autofluorescent protein tagging and expression of multiple genes in plants. Plant Mol. Biol. 57, 503–16 (2005).

44. Liu, Q., Li, M. Z., Leibham, D., Cortez, D. & Elledge, S. J. The univector plasmid-fusion system, a method for rapid construction of recombinant DNA without restriction enzymes. Curr. Biol. 8, 1300–9 (1998).

45. Schneider, C. A., Rasband, W. S. & Eliceiri, K. W. NIH Image to ImageJ: 25 years of image analysis. Nat. Methods 9, 671–5 (2012).

